# Genetic and behavioural repertoires support olfactory communication in wild capuchin monkeys

**DOI:** 10.64898/2026.09.08.750240

**Authors:** MS Akhtar, NK Kulick, AC Poirier, H Hamou, CF Galán-Plana, S Romero Morales, S Cheves Hernandez, GW Benson, C Chaves-Cordero, R Hernández-Rojas, V Villero, M Crofoot, T Bergman, JD Orkin, BJ Barrett, J Tung, KM Jack, AD Melin

**Author notes:** denotes co-first authorship (MS Akhtar, NK Kulick). Author for correspondence: Muhammad Shoaib Akhtar. Amanda D Melin.

## Abstract

Olfaction is a key sense for many mammals, but its role in the social lives of anthropoid primates (monkeys, apes, humans) is poorly understood. Studies of chemosensory receptor genes and olfactory behaviours can provide insight into the evolution and function of olfaction across social contexts. Vomeronasal receptor (*VR*) genes are especially relevant to social behaviour as they are specialized for the detection of pheromones and other cues involved in social signaling, reproductive behaviour, and individual recognition in mammals. Here, we examined the population genetics and gene repertoires of *VR* genes in wild white-faced capuchin monkeys (*Cebus imitator*) from Costa Rica and Panama using DNA sourced through both host tissues and through noninvasive fluorescence-activated cell sorting of fecal samples (fecalFACS). We compared *VR* diversity relative to putatively neutral reference loci (NRs) to investigate evidence of selection and potential effects of demography. To complement the genetic analysis, we also conducted a detailed observational study of capuchin olfactory behaviours. Combined, we find evidence of intact *VRs* subject to evolutionary constraint in capuchins, providing novel evidence that members of this gene family are maintained by evolutionary constraint in wild monkeys. We further document a rich and varied ethogram of olfactory behaviours that have clear potential for communication. Overall, our study provides compelling evidence for a role of olfaction in the complex social lives of capuchin monkeys, motivating future research on the contributions of olfaction to social communication and evolution in primates and other mammals.

## Introduction

Olfaction is a pillar of social communication in mammals, assisting in diverse functions including territory marking (rodents: Beauchamp & Yamazaki, 2003; Hurst & Beynon, 2004; canids: Apps et al., 2012; Cafazzo et al., 2012; Sillero-Zubiri & Macdonald, 1998; felids: Allen et al., 2017; Soso & Koziel, 2016; Zaman et al., 2024), kin identification (mongooses: Jordan et al., 2010; Mitchell et al., 2018; bats: Liang et al., 2021; Safi & Kerth, 2003), communicating emotional state (elephants: Rasmussen & Krishnamurthy, 2000; humans: Calvi et al., 2020; Roberts et al., 2022), and signalling fertility (otters: Kean et al., 2011; giant pandas: Swaisgood et al., 2002; Zhou et al., 2019). Olfactory communication through volatile and semi-volatile signals has the greatest breadth and time-depth of any sensory modality: it can be useful over distances of several kilometers or more and persist for up to several weeks (Dominy et al., 2004; Higham & Hebets, 2013; Wyatt, 2014). While olfactory communication has been studied in depth for some species, the importance of olfaction in social contexts is less understood for many primates (Drea, 2020; Heymann, 2006b). This lack of knowledge is especially pronounced for the anthropoid primates (monkeys, apes, humans), which is perhaps surprising given how intensively studied this diverse radiation of mammals has otherwise been (Rowe & Myers, 2016).

The paucity of studies on olfactory communication in anthropoids stems in part from a historical emphasis on the loss of olfactory adaptations and gain of visual adaptations in this radiation of primates (Gilad et al., 2004; Kawamura & Melin, 2017; Laska et al., 2005; Liman & Innan, 2003; Niimura et al., 2018). The anthropoid visual system has expanded in size and complexity with numerous derived features, including increased eye size, binocularity and, in the diurnal anthropoids (monkeys and apes), a cone-dense fovea (Gilad et al., 2004; Hiramatsu et al., 2017). These morphological characteristics, together with the bright colouration of many monkeys, has inspired a large body of research that has established a central role of vision in primate social communication (e.g., Higham et al., 2008, 2021; Moreira et al., 2019; Setchell et al., 2006; Setchell & Wickings, 2005). Conversely, features of the olfactory system of anthropoids are relatively diminished: they have lost a wet rhinarium, have fewer genes underlying olfactory perception than many other mammals, and have relatively small olfactory bulbs (Niimura et al., 2018; Smith et al., 2007; Smith & Rossie, 2006). Despite these changes, monkeys retain many key features, molecular and anatomical, relevant to social olfaction. While sparse, available evidence suggests the sense of smell may be involved in diverse behaviours spanning reproduction, social bonding, dominance interactions, and territoriality in multiple primate taxa, including monkeys (Charpentier et al., 2016; Chi et al., 2025; Drea, 2015; Hiramatsu et al., 2017; Poirier & Melin, 2024; Smith et al., 2001; Veilleux et al., 2022).

The number and functional integrity of chemosensory receptor genes have frequently been called upon to provide insight into the evolution of olfaction. There are two main classes of olfaction-related chemosensory receptors: olfactory receptors (ORs, coded by *OR* genes) and vomeronasal receptors (VRs, coded by *VR* genes). The *OR* genes encode the receptors in the olfactory epithelium of the nose. They are typically stimulated by fully volatile odorants, which induce a neural signaling pathway that facilitates perception. *OR*s are diverse and respond to a wide variety of ecological as well as putative social olfactory signals (Buck & Axel, 1991; Issel-Tarver & Rine, 1997; Malnic et al., 1999; Mohrhardt et al., 2018). The *VRs* are part of the vomeronasal organ (VNO) system which is present in numerous species across the animal kingdom and typically specialized for social signaling and olfactory communication among conspecifics (D’Aniello et al., 2017; Silva & Antunes, 2017). The composition of *VR* genes is diverse and varied in many mammalian species, including strepsirrhine primates (lemurs and lorises), which have a relatively large VNO (Smith et al., 2015). However, the VNO has been reduced in anthropoids and seemingly completely lost in some catarrhine primates (monkeys and apes in Africa, Asia, and Europe; Bhatnagar & Smith, 2014; Yoder & Larsen, 2014; but see Charpentier et al., 2013; Smith et al., 2001). In contrast, some evidence suggests that functional *VR* genes persist in some platyrrhine primates (monkeys in the Americas; Moriya-Ito et al., 2018; Smith et al., 2011), but both the empirical research and data in this area are sparse.

In addition to genetic, morphological, and physiological adaptations to support olfactory perception, odour-based communication requires multiple participants: one to emit signals or cues (the signaller) and one or more others to perceive them (the receiver(s)). In mammalian olfactory communication, scent marking – the rubbing of odorous secretions from specialized scent glands and/or the distribution of excretions, including urine and feces – has evolved repeatedly in diverse lineages as a signalling behaviour (Brown & Macdonald, 1985; Johnson, 1973). Additionally, olfactory signalling is possible through more passive deposition of body odours in the environment, for example through resting in a certain area (Albone, 1984; Wyatt, 2014). On the receiver’s side, scent perception may occur passively through inhalation or actively through specialized behaviours that enhance the uptake of volatile and semivolatile odorants, such as sniffing or performing the flehmen response, which entails curling up the lips while inhaling deeply (Schneider, 1930). Licking and tasting can also contribute to olfactory perception (Charpentier et al., 2013; Henkel & Setchell, 2018; Smith et al., 1997). The extent and manner in which individuals engage in these signalling and receiving pathways is expected to reflect their use of olfactory communication. Accordingly, characterizing putative olfactory behaviours, and studying if and how natural selection acts on chemosensory genes in natural systems, should provide additional insight into the social importance of olfaction (Baum & Cherry, 2015; Delbarco-Trillo et al., 2011; Drea, 2015; Heymann, 2006a; Hiramatsu et al., 2009).

In this study, we investigated the genetic and behavioural potential for social olfactory signaling in a platyrrhine primate species, the white-faced capuchin monkey (*Cebus imitator*). Specifically, we paired population genetic analysis of putative *VR* genes across multiple populations in Central America with detailed observation of olfactory-related behaviours in a wild population at our primary study site in northwest Costa Rica. We pursued two aims: 1) to seek evidence for the presence, functional integrity, and selective pressures acting on *VR* genes, and 2) to assess the behavioural evidence that olfaction is used regularly in socially-relevant contexts in this species. Under the hypothesis that mechanisms supporting olfactory communication are maintained by natural selection, we predicted that we would find evidence of functional *VR* genes carrying a signature of evolutionary constraint, and that capuchins would show chemosensory behaviours (sniffing, tasting) in socially-relevant contexts.For Aim 1, we **1a)** generated and searched white-faced capuchin genome sequences for the presence and intact structure of *VR* genes, and **1b)** analyzed population genetic diversity of these genes and putatively neutral loci to evaluate the operation of evolutionary constraint maintaining *VR* genes in white-faced capuchins. In Aim 2, we **2a)** developed a comprehensive and detailed ethogram of olfactory behaviours, and **2b)** reported frequencies, contexts, and examples of putative olfactory social behaviours. We also evaluated whether olfactory behaviours of interest vary seasonally, as capuchins in the study population are moderately seasonal breeders and most births, male emigrations, and alpha male replacements occur in the dry season (Brasington et al., 2022; Fedigan, 2003). Our goal was to provide a comprehensive study in a well-habituated population, and to inspire additional future research on the mosaic of sensory evolution and communication in mammals, including those that have not historically been considered olfactory specialists.

## Materials and Methods

### Study species

White-faced capuchins (*Cebus imitator*) are medium-sized, omnivorous and highly social platyrrhine monkeys endemic to Central America (Fragaszy et al., 2004). They reside in female philopatric, multimale-multifemale groups of 5-40 individuals (Jacobson et al., 2025; Melin et al., 2020), with an adult sex ratio of around two females per male (Fedigan & Jack, 2011). Olfactory communication in white-faced capuchins may underlie diverse intragroup and intergroup social interactions. We focused our behavioural study on males as they: 1) are more likely than females to perform putative signalling (urine washing) and receiving (sniffing) behaviours (Campos et al., 2007; personal observations, KMJ and NKK), 2) are more engaged in territorial interactions (Perry, 1996b), and 3) show bimaturism in adult body phenotypes (Chapoy et al., 2025). In other mammalian species, the emergence of these phenotypes are influenced by olfactory signalling (e.g., house mice; Hurst et al., 2001), raising the possibility that they may also do so here.

Males typically disperse from their natal group around the age of 4-5 years, and continue to disperse between groups every ∼3-4 years throughout adulthood (Jack & Fedigan, 2004b, 2004a). While females maintain linear dominance hierarchies within groups (Perry, 1996a), males exhibit two distinct dominance ranks: a single alpha male position per group, characterized by increased centrality, vigilance, and a more robust morphology, and a subordinate position occupied by the rest of the group’s adult males (Chapoy et al., 2025; Jack & Fedigan, 2018). In our primary study population, in Sector Santa Rosa (SSR), Área de Conservación Guanacaste, Guanacaste, Costa Rica, alpha male replacements occur every ∼3 years, typically during the dry season, and represent a time of increased intragroup aggression as males compete for the alpha position (Petersdorf et al., 2025; Teichroeb & Jack, 2017). At any given time, male reproductive skew is high, with alpha males siring ∼80% of offspring (Jack & Fedigan, 2006; Muniz et al., 2010; Wikberg et al., 2017). Females have visually cryptic ovulation (Fedigan & Jack, 2013), lacking the obvious sexual swellings seen in some catarrhine primates, and most conceptions occur during the rainy season (Brasington et al., 2022). Within-group aggression among males and among females is relatively rare during times of hierarchy and group stability (Bernstein, 1976; Fedigan, 1993).

White-faced capuchins have rich social lives, with individuals maintaining close social contact throughout the day while foraging, feeding, traveling, resting, grooming, playing, or engaging in other social affiliative behaviours (e.g., social rubbing of plants on each others’ fur; (Fragaszy et al., 2004; Perry, 2012; Perry et al., 2003). Females maintain close bonds with their female family members throughout their lives (Perry, 1996a), and males often also maintain lifelong affiliative relationships with other males (kin and nonkin) as they disperse in parallel from their natal group and between subsequent groups (Jack & Fedigan, 2004b; Perry, 1998). This species is moderately territorial. Their yearly home ranges in tropical dry forest average 1.82 km² (range = 0.35-7.29 km², Jacobson et al., 2024) to 1.98 km² (SD = 0.59 km², Campos et al., 2014), and while home ranges often greatly overlap with those of other groups (Jacobson et al., 2026), capuchins are highly aggressive towards other groups and extra-group males. When intergroup encounters occur, adult males will pursue and fight with the males of the neighboring group. To reunite with the rest of their group following an encounter, males will search the environment and often vocalize with loud lost calls which are typically answered by group members (Fragaszy et al., 2004; Perry, 1998; Schoof & Jack, 2013). If olfactory communication is socially important in white-faced capuchins, olfactory behaviours would be expected under such competitive, sexual, and affiliative circumstances.

### Study sites

#### Sites for genomic sampling

To investigate the complement and diversity of putative *VR* genes in white-faced capuchins, we first generated genome-wide resequencing data from four ecologically distinct sites, representing major forest biomes in Costa Rica and Panama (Figure 1). The sampled capuchin monkeys (n=95 individuals) included infants, juvenile, and adults, comprising 46 males, 39 females and 10 monkeys for which sex was unknown (S Table 1).

**Figure 1:**
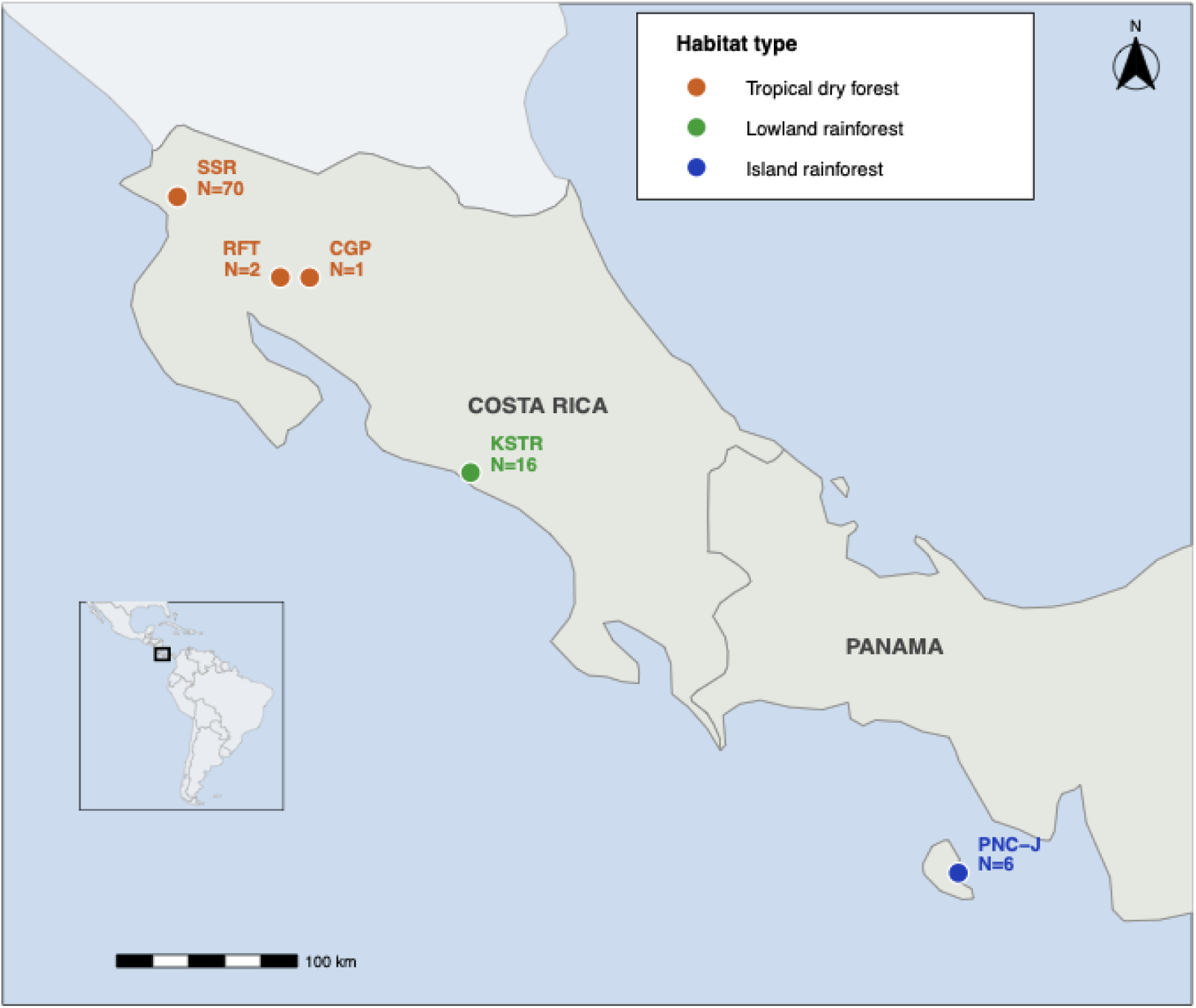
Sampling locations of wild capuchin monkeys (*Cebus imitator*).

Seventy of the monkeys were sampled from our primary study site in SSR, Área de Conservación Guanacaste, Guanacaste, Costa Rica (Supplementary Table 1). SSR is a largely secondary regrowth tropical dry forest environment that experiences intense seasonality, with ∼6 months of daily precipitation during the rainy season (late May/early June through November; average 100 mm/month) followed by ∼6 months during which nearly no rain falls and a majority of the drought-deciduous trees in the forest lose their leaves (late November/early December into May; Campos et al., 2014; Hogan & Melin, 2018). DNA from monkeys in SSR was primarily collected noninvasively from fecal samples, although we sampled tissues from seven monkeys who died of natural causes during our study. Additionally, DNA from urine was sampled from two SSR monkeys. Outside of SSR, two capuchin samples were collected from Reserva Forestal Taboga (RFT) in a tropical dry forest near SSR in Costa Rica, surrounded by agricultural lands, and one sample came from a monkey who originated near Canas in Guanacaste province (CGP) and coded as SSR in this study. We also sampled blood or tissues from 16 capuchins living in dense lowland rainforest typical of the Pacific coastal zone in the Quepos and Manuel Antonio region. These samples were collected by veterinarians at the KSTR Wildlife Rescue Center (KSTR), where the primates were undergoing treatment. This habitat is characterized by high annual rainfall, evergreen vegetation, and humid coastal conditions (Kappelle, 2016). Finally, DNA samples from a Panamanian population of the same capuchin species (n=6) were obtained from collecting feces from capuchins living on Jicarón island (PNC-J). Jicarón Island, part of Coiba National Park, is a humid tropical lowland rainforest, with high precipitation, evergreen forest cover, and minimal human disturbance (Barrett et al., 2018; Castroviejo & Velayos, 1997). Together, these four sites capture a broad environmental gradient, including tropical dry forest, wet lowland rainforest, and island systems with contrasting moisture regimes, providing ecological context for investigating *VR* diversity across capuchin monkey populations (Figure 1).

#### Site of behavioural study

For our second aim focused on olfactory communication, we studied the behaviour of capuchins in our long-term study population at Sector Santa Rosa (SSR), Área de Conservación Guanacaste, Guanacaste, Costa Rica, described above. The SSR capuchins have been studied nearly continuously since 1983, resulting in comprehensive longitudinal datasets on ecology, demography, and the life history and behaviours of most individuals (Melin et al., 2020). During the dry season there is excellent visibility in the forest (Campos et al., 2024), facilitating detailed descriptive behavioural observations of sensory behaviours (Melin et al., 2022)

### Genomic data generation and analysis

#### DNA extraction, library preparation, and sequencing

We generated genome resequencing data from 101 biological samples (73 fecal samples, 12 blood samples, 4 ear samples, 7 kidney samples, 2 testes samples, 2 urine samples, and 1 unknown origin sample), representing 95 unique individuals. Blood and tissue samples typically generate high-quality DNA extracts that overwhelmingly represent the DNA of the individual from whom they were collected. We therefore extracted DNA from these 28 samples (all except fecal samples) using standard commercial kits: Qiagen DNEasy Blood and Tissue kit (69582) for the blood samples, and the Machery Nagel Nucleospin Tissue kit (MN-740952), for the tissue samples, following the manufacturer’s instructions.

In contrast, direct DNA extracts from noninvasively collected fecal and urine samples, the only sample type possible to collect for some populations are typically low quality and often overwhelmingly microbial (Chiou & Bergey, 2018; Snyder-Mackler et al., 2016; Tang et al., 2025; van der Valk et al., 2017). To overcome this and isolate DNA suitable for high-throughput sequencing, we sorted monkey cells from the fecal or urine matrix using a fluorescence activated cell sorting protocol (fecalFACS) at Flow Cytometry Core Facility of the University of Calgary previously described in Orkin et al. (2021), followed by DNA extraction from sorted, host-enriched cells using the QIAamp DNA Mini Kit (56304) or Thermo Fisher PicoPure DNA Extraction Kit (KIT0103). FecalFACS-enriched samples were highly enriched for endogenous host DNA relative to DNA extracts obtained directly from fecal material.

For all sample types, we built DNA-seq libraries with the NEB Next Ultra 2, xGen cfDNA & FFPE DNA, or xGen™ ssDNA & Low-Input DNA Library Preparation Kits following the manufacturers’ protocols and sequenced the resulting libraries at either the University of Calgary Genome Sequencing Core or the Core Unit of the Max Planck Institute for Evolutionary Anthropology (S Table 1).

#### Sequencing read processing and quality control

Our primary goal was to identify and extract putative *VR* gene sequences from the larger resequencing data set to study the *VR* complement and its genetic diversity in white-faced capuchins. To do so, we first subjected the resequencing data set to standard mapping, variant calling, and quality control pipelines, including removal of adapter sequence, trimming of low-quality bases, and removal of potential human contamination (Supplementary Methods). The remaining reads were mapped to the *Cebus imitator 1.0* genome assembly with *Bowtie2* (Langmead & Salzberg, 2012). We removed PCR duplicates and reads with a mapping quality score of less than 10, merged BAM files from the same individual, and performed variant calling using GATK HaplotypeCaller following GATK Best Practices (Poplin et al., 2018; Van der Auwera et al., 2013). The final dataset included 80 individuals, while rest were excluded as part of minimal depth and quality filters.

We then focused specifically on *VR* genes and complementary putatively neutral regions for comparison (see below and Table 1 and Table S2). Following the variant calling pipeline, we annotated the variants using snpEff (Cingolani et al., 2012). Finally, we built a custom snpEff database for the *Cebus imitator* 1.0 genome assembly using annotations provided in the assembly’s general feature format (GFF).

**Table 1.**
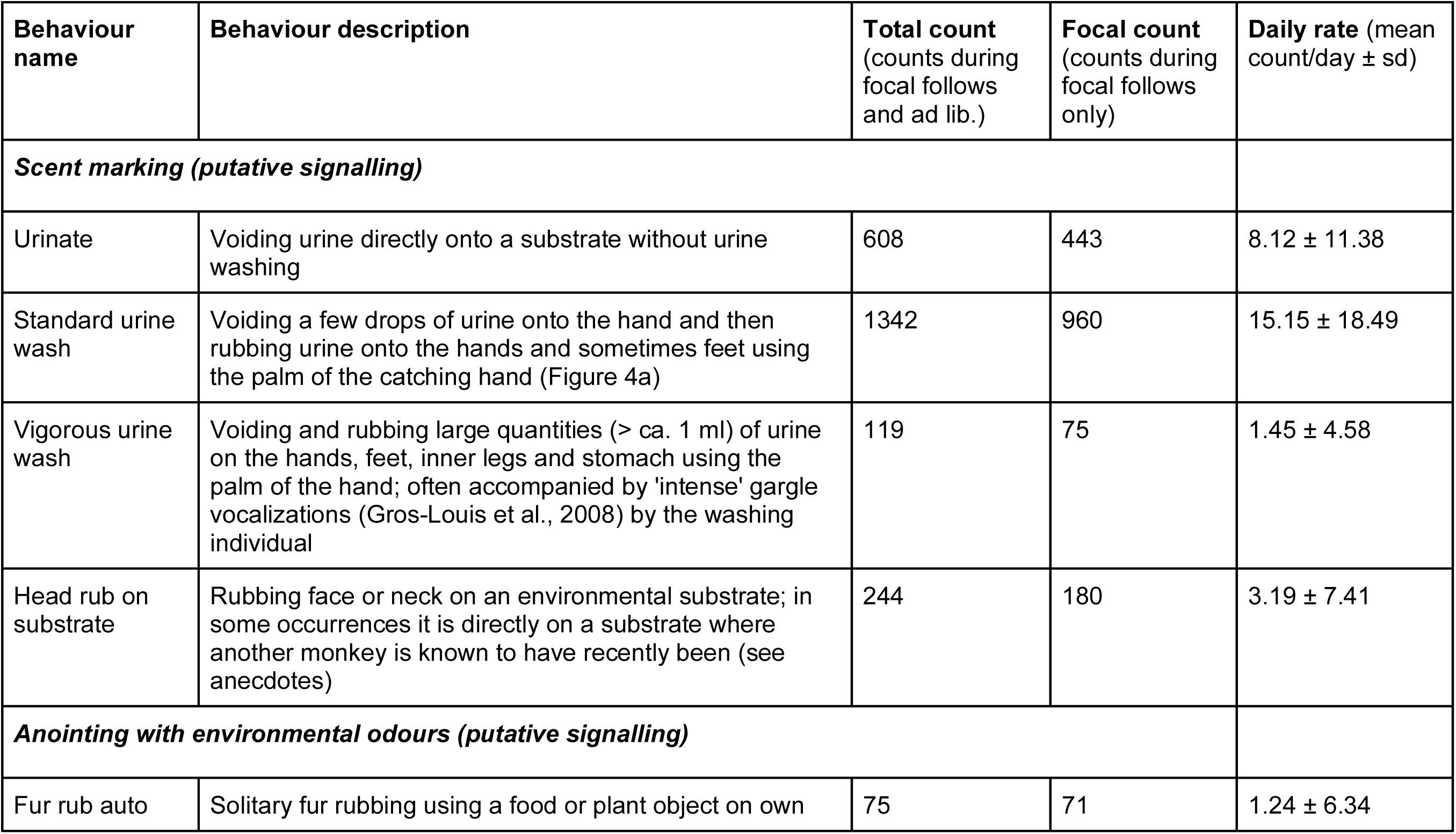
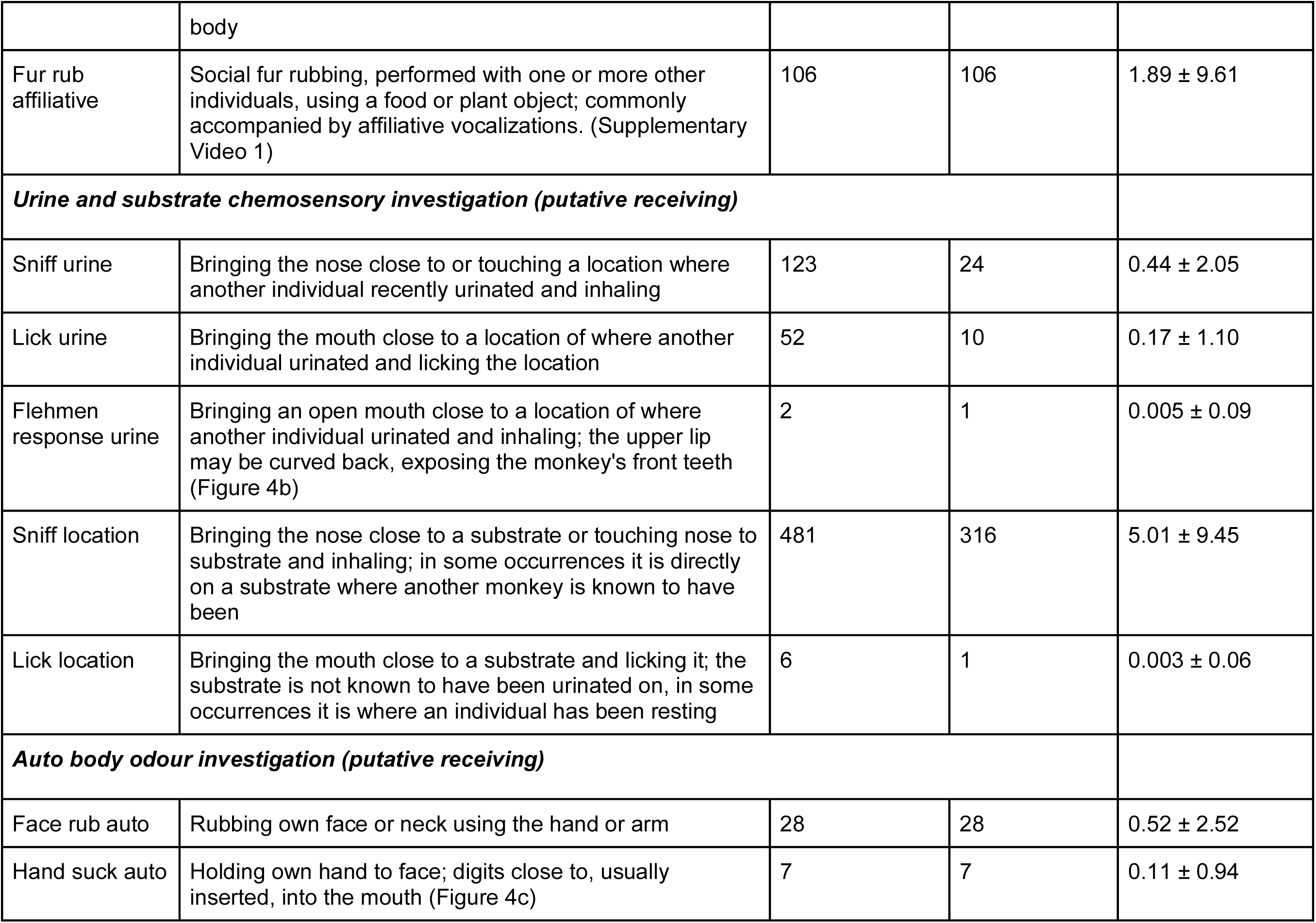
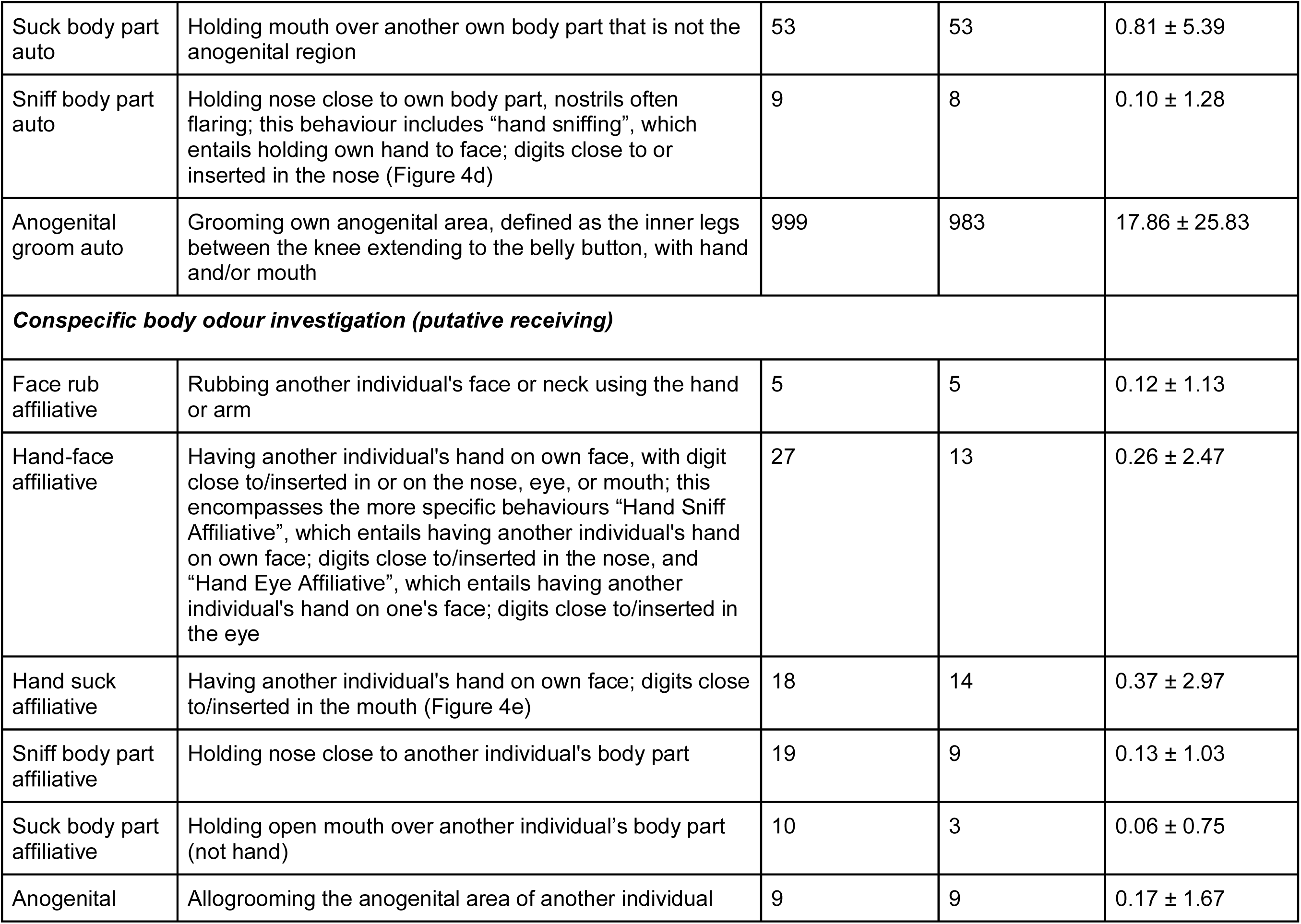
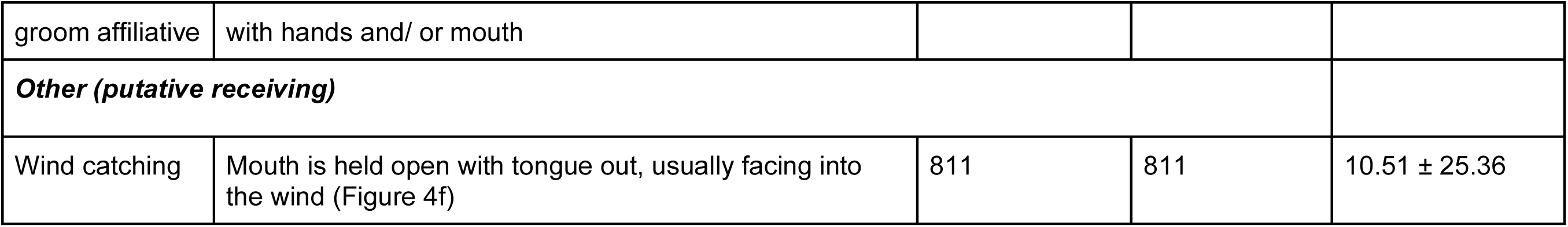
Ethogram of putative socially-informative olfactory behaviours, their descriptions, and their counts and daily frequencies. Total count represents the total number of times each behaviour was observed during focal follows and ad libitum (3,022.75 contact hours) and focal count represents the number of times each behaviour was observed during focal follows (789.95 focal hours). Daily behavioural rates (count/day) were calculated as hourly behavioural incidence rates based on focal follow durations, which we then used to estimate daily rates assuming 12 waking hours per day. In Supplementary Video 1 we present examples of many of these behaviours

#### Identification of putative VR and NR sequences in the white-faced capuchin genome

To define *VRs*, we used the vomeronasal receptor (*VR*) coordinates that were tentatively identified in the *Cebus imitator 1.0* genome assembly by Orkin et al. (2021). In that study, the authors identified seven putative *VRs*, four of which were annotated as intact and three as pseudogenes. In addition, we used a set of sequences for *VRs* from all possible primate genomes including black-capped squirrel monkey, bonobo, chinese tree shrew, common chimpanzee, common marmoset, crab-eating macaque, golden snub-nosed monkey, gray mouse lemur, green monkey, human, northern greater galago, northern white-cheeked gibbon, Philippine tarsier, rhesus macaque, sumatran orangutan, and western gorilla (Moriya-Ito et al., 2018; Hohenbrink et al., 2017; Hunnicutt et al., 2020). We then aligned these *VR* sequences against *Cebus imitator 1.0* genome assembly, and identified putative *VR* genes (For more details, S Methods 2). We applied the Tamura 3-parameter model to construct a phylogenetic tree using Maximum-Likelihood method and 1000 bootstrap replicates to identify paralogous genes. We extracted the sequences of these *VRs* to validate the intact/pseudogene classifications. We used EMBOSS getorf (Rice et al., 2000) to confirm that the open reading frame of each *VR* gene was at least 250 amino acids long, the length necessary to form a seven-transmembrane structure required by functional G-protein coupled receptors. We then used DeepTMHMM, a neural network-based approach (Hallgren et al., 2022), to identify open reading frames of 250 amino acids or longer that are capable of making a seven-transmembrane structure. If a *VR* gene met both criteria, then we considered it intact; otherwise, it was labelled as disrupted. For the identified *VR* genes with lengths between 500 and 750 nucleotides were labelled as truncated. We note that our analyses are based solely on genomic assessment and our results do not confirm the genes’ functional status. For this reason, we use “intact” and “disrupted” gene terminologies rather than “functional” and “pseudogenes.” Following these criteria, we identified 12 intact, 28 disrupted, and seven truncated *VR* genes.

After identifying the *VR* coordinates, we identified a representative subset of putatively neutrally evolving reference regions (NRs) in the *Cebus imitator* 1.0 genome assembly. NRs are large, repeat-free intergenic regions that are not part of any coding region and at least 0.2 centimorgan away from the nearest gene in autosomes (based on Neutral Region Explorer (Arbiza et al., 2012). We aligned NRs previously reported for the hg19 human reference genome to *Cebus imitator 1.0* (Akhtar et al., 2022) to identify 64 capuchin NRs with similarity of more than 75%, which reflects the expected level of neutral sequence divergence between non-coding ortholog sequences between humans and capuchins. Alignment settings used were the same with the ones used in *VR* identification and are given in S Methods 2. Recent studies have used the same NR dataset, aligned to several anthropoid primate genomes, to confirm presence of NR orthologs (Hou et al., 2024; Ji et al., 2025). We selected only the highest similarity sequence for each query, with a minimal length of 1 kb, after confirming that this sequence was not part of any annotated protein-coding region within the capuchin genome.

#### Population genetic analyses

To understand patterns of variation and potential signatures of selection in capuchin *VRs*, we analyzed allele frequencies, nucleotide diversity (π), and Tajima’s D in both *VRs* and NRs. To test our prediction that intact *VR* genes are functionally constrained, we used Tajima’s D to test for departures from neutrality and haplotype frequency estimation to test for limited haplotype diversity. Tajima’s D is calculated based on the difference in the number of pairwise differences (π) and the number of segregating sites (s), scaled by sample size, in a genomic region (Tajima, 1989), and was calculated using the vcf2tajima tool (Akhtar & Kawamura, 2024). We calculated allele frequencies and π per site in our sample using VCFTools (Danecek et al., 2011).

To further investigate genetic diversity at capuchin *VR* genes, we phased our final dataset of 80 capuchin genomes to identify haplotype sequences of 12 intact VRs and the 64 NRs using BEAGLE (Browning et al., 2021), which infers haplotypes based on localized linkage disequilibrium patterns. To evaluate whether each haplotype sequence encoded an intact *VR*, they were assessed to confirm an open read frame and the potential to make a seven-transmembrane structure. Phased haplotypes were subsequently used for haplotype frequency estimation and a haplotype network was constructed using the *popart* minimum spanning network (Leigh et al., 2015). Predominance of a single haplotype, along with several rare, closely related haplotypes is a signature of natural selection and functional constraint. While we expected such a signal for *VR*s, under the hypothesis that they are maintained by natural selection, we expected NRs to depart from this pattern.

### Behavioural methods

#### Field behavioural observations

To generate a detailed ethogram of putative chemosensory behaviours, we performed focal animal follows (Altmann, 1974) on adult male wild capuchins 7 years and older (N=47) in 6 social groups from March–May 2021 and January 2022–June 2024. While in contact with a study group (6:00-18:00), we observed and recorded all occurrences of behaviours putatively relevant to the perception of social olfactory cues and signals (sniffing/ licking) and behaviours that were putatively involved in generating cues and signals (e.g., rubbing the face or neck on substrates, urine washing). To be included as a behaviour in the formal ethogram, we required the behaviour to be consistently and reliably identified by each of our team members, such that two researchers observing the same monkey would record the same behaviour code with regularity (>95% of the time) during focal animal follows. Focals were performed on 382 days, with an average of 9.88 performed each day (sd 6.03 focals). During 2021, focals were designed to optimize continuous observation time of individuals (mean duration 97.32 min, sd 98.73 min). From 2022-2024, our focal durations were changed to the more standard duration of 10 minutes (mean duration 10.5 min, sd 2.26 min). Additionally, we recorded all observed olfactory behaviours *ad libitum* outside of focal follows on all adult males and females in the study groups. Using these data, we report the frequencies of these behaviours and describe representative cases regarding their social and ecological contexts.

#### Statistical analyses

To assess patterns in the seasonality of the 23 behaviours in our ethogram (Table 2), we ran a generalized linear mixed effects model (GLMM) for each behaviour fit with a negative binomial distribution and log link function (glmmTMB R package; McGillycuddy et al., 2025).

Each model tested the effect of rainfall seasonality, using average daily precipitation values for the 30 days prior to each sampling date (Orkin et al., 2021; Poirier et al., 2025), on daily behavioural counts. We included an offset based on the daily log-transformed duration of focal sampling to account for differences in focal follow protocols during the study period. We assessed the contribution of seasonality on model fit for each behaviour using a likelihood ratio test.

All reported models met model assumptions and showed no indication of zero inflation or overdispersion (assessed with the DHARMa R package; Hartig, 2025). The models for five behaviours failed to meet model expectations, likely due to their small sample sizes, and were excluded, resulting in tests for 18 behaviours in the ethogram. We evaluated predictor significance at the alpha=0.05 level.

For each behaviour, seasonal occurrence was visualized using circular histograms (rose plots). Following Melin et al. (2014), the mean vector was calculated from the day of year, converted to radians, and weighted by the number of observations per 12-hour day. Mean vector length (r), which quantifies the concentration of of observations around the annual cycle, was calculated as: √((sum(count per 12 hour day × cos(day of year in radians)) / sum(count per 12 hour day))² + (sum(count per 12 hour day × sin(day of year in radians)) / sum(count per 12 hour day))²). All behavioural analyses were conducted in R v4.5.0 (R Core Team, 2025).

## Results

### The vomeronasal gene receptor repertoire in white-faced capuchins

Twelve *VR* genes met our definition of intactness, i.e., an open reading frame of at least 250 amino acids and potential to make a seven transmembrane structure. Another 28 *VR* genes identified in the genome did not meet our criteria for intactness and exhibited evidence of disruption, including two *VRs* that had an open reading frame of at least 250 amino acids but were modeled to be unable to make a seven transmembrane structure. Additionally, we identified seven *VR*s with nucleotide sequence length between 500 and 749 nucleotides and classified them as truncated. Although truncated *VR* gene sequences are shorter than this threshold in current assembly, assembly artifacts may contribute to truncations. Any *VR* gene-like sequences which were shorter than 500 nucleotides are not included in gene repertoire classification. Out of the 48 identified *VR*s, 47 belonged to the type 1 vomeronasal receptor family (*V1Rs*), whereas one belonged to the type 2 family (*V2R*s). The only *V2R* gene was *V2R1P*, which was found to be a disrupted gene. The *V2R1P* was previously identified by Orkin et al. (2021) and we have looked for its coding sequence which was only 169 nucleotides long and thus identified as disrupted. The phylogenetic tree of all capuchin *V1Rs* showing paralogous relationship is given as S Figure 2. All *V1R* paralogous gene groups were identified and named based on their phylogenetic relationship. The phylogenetic tree also shows intact and disrupted *V1R* genes interspersed with each other which reflects multiple independent disruption events. A list of all these genes, genomic coordinates, their status as intact, disrupted or truncated, and orthologous gene information is given in S Table 2.

In addition to *VR* gene repertoire, we identified 64 NR sequences with lengths between 1014 and 6105 nucleotides in capuchin genome assembly and a list of these NRs with their genomic coordinates is given in S Table 3. Before variant calling, we calculated mean depth per *VR* gene or NR region for each individual which was 13.66 ± 10.03x for intact *VRs*, 13.60 ± 10.46x for disrupted *VRs*, and 13.36 ± 10.47x for NRs.

### Variation in the VR repertoire across populations and individuals

We investigated inter-individual variation in *VR* repertoires for each of the 12 intact, 28 disrupted and seven truncated *VR* genes identified in the white-faced capuchin genome assembly. We used the 64 NR sequences as a comparator in all analyses. To investigate inter-individual variation, we focused on the 80 individuals (of 95 originally targeted for sequencing; S Table 4) where we achieved moderate to high sequencing depth.

We identified 223 variants in the 12 intact *VRs*, including 112 single nucleotide variants (SNVs), 76 insertions, and 35 deletions. In the 28 disrupted or truncated *VRs*, we identified 614 variants, including 380 SNVs, 179 insertions, and 75 deletions (Figure 2A). Among disrupted *VR* gene variants, 96 SNVs, 64 insertions, and 22 deletions were observed in *V2R* gene, and rest were in *V1Rs*. We also called 3267 variants (1875 SNVs, 928 insertions and 464 deletions) in the 64 NR loci (Figure 2A).Mean π for segregating sites was 0.032 ± 0.075 in intact *VRs*, 0.068 ± 0.130 in disrupted and truncated *VRs*, and 0.067 ± 0.129 in NRs, consistent with the hypothesis of greater evolutionary constraint in intact *VRs* versus disrupted *VRs* or NRs. VCF files of both *VRs* and NRs are given online in Zenodo Repository.

**Figure 2.**
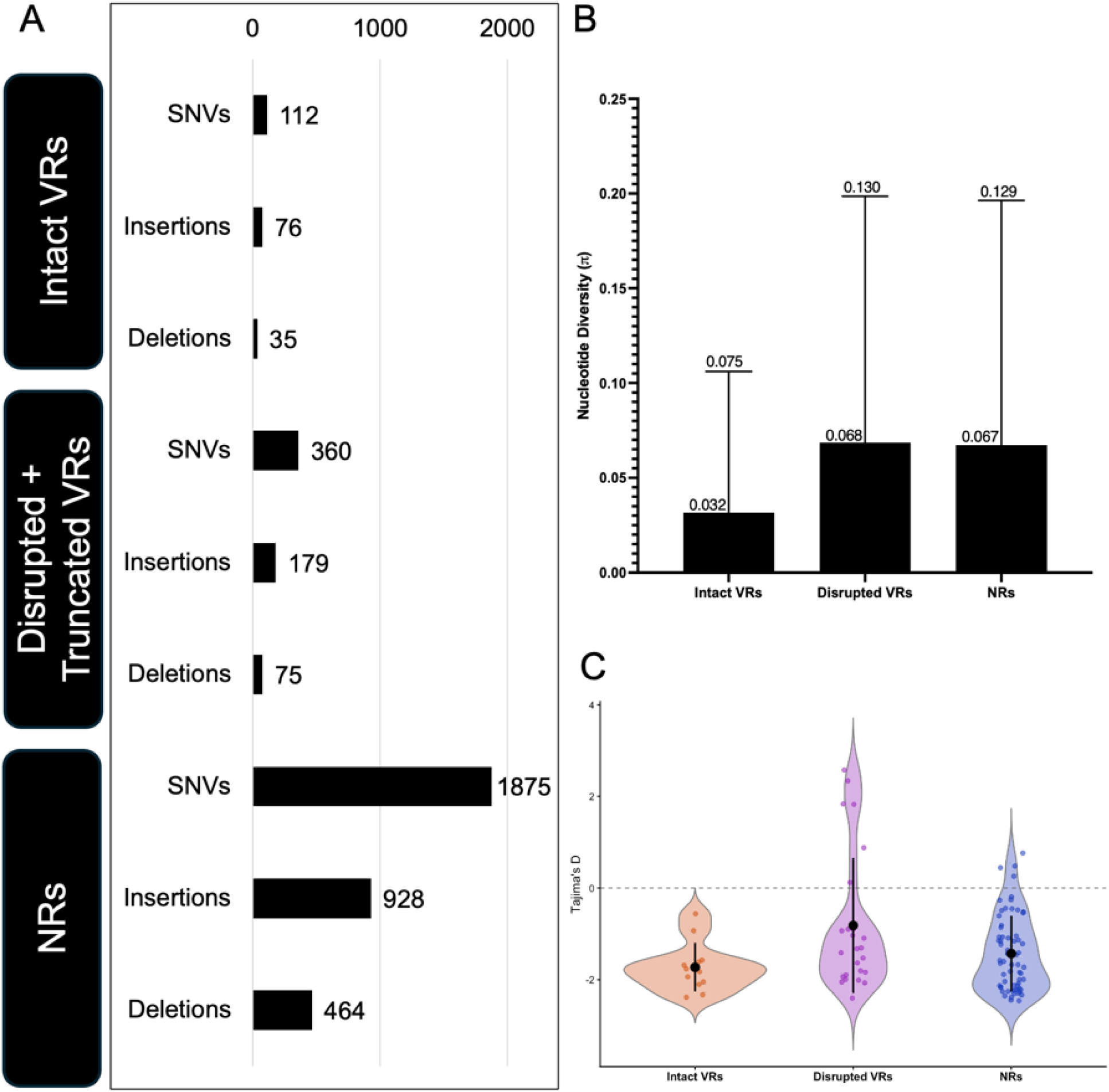
*VR* repertoire in wild capuchin monkeys. A: Number of SNVs and insertions/deletions (relative to the *Cebus imitator 1.0* reference genome) in intact, disrupted and truncated VRs, and NRs in n=80 white-faced capuchins from Costa Rica and Panama. B: Distribution of π for segregating sites in intact, disrupted and truncated *VRs*, and NRs in the same data set. C. Tajima’s D of intact *VRs,* disrupted and truncated *VRs* and NRs. Plot of Tajima’s D for each intact *VR* gene and NR region in the current study, where each dot shows Tajima’s D for one locus and vertical line within violin shows mean and standard deviation within each group.

### Signature of evolutionary constraint

We investigated possible evolutionary constraints in intact *VR* genes by comparing Tajima’s D for intact and disrupted *VR* genes to Tajima’s D for NRs. All intact *VR* genes had negative Tajima’s D values (mean -1.73 ± 0.53). In contrast, Tajima’s D was highly variable for disrupted *VRs* and NRs, with both negative and positive values and a mean relatively closer to 0 (disrupted *VRs*: mean -0.88 ± 1.48; NRs: mean -1.43 ± 0.83 Figure 2C). While mean Tajima’s D for intact *VRs* was lower than mean Tajima’s D for NRs, mean Tajima’s D for disrupted *VRs* was higher than both. This lower Tajima’s D observation at intact *VR* loci is consistent with possible evolutionary constraints, in contrast to the other regions examined. In contrast, the higher Tajima’s D observation at disrupted *VRs* is consistent with possible relaxed evolutionary constraint following loss of function allowing variants to accumulate.

### VR haplotypes

We identified unique haplotypes for all 12 intact *VRs* and 64 NRs. We showed distribution of these haplotypes as a minimum spanning network within study individuals. For all intact *VRs*, there is only one haplotype common across all study individuals followed by the existence of some low frequency haplotypes. This common haplotype is intact in all cases, but some of the low frequency haplotypes are disrupted in each *VR* gene. In case, an intermediate frequency haplotype was observed in any *VR* gene along with most common haplotype then this haplotype was also intact. This pattern is shown in Figure 3 (A-F; and S Figure 3) as the strong skew in haplotype frequencies towards the most common and intact haplotypes observable in a haplotype minimum spanning network dominated by a large node. This pattern contrasts with the more distributed variation observable for NRs (Figure 3G-L and S Figure 4), where no single haplotype dominantes, and is consistent with greater evolutionary constraint at the intact *VRs* than at neutrally evolving regions.

**Figure 3:**
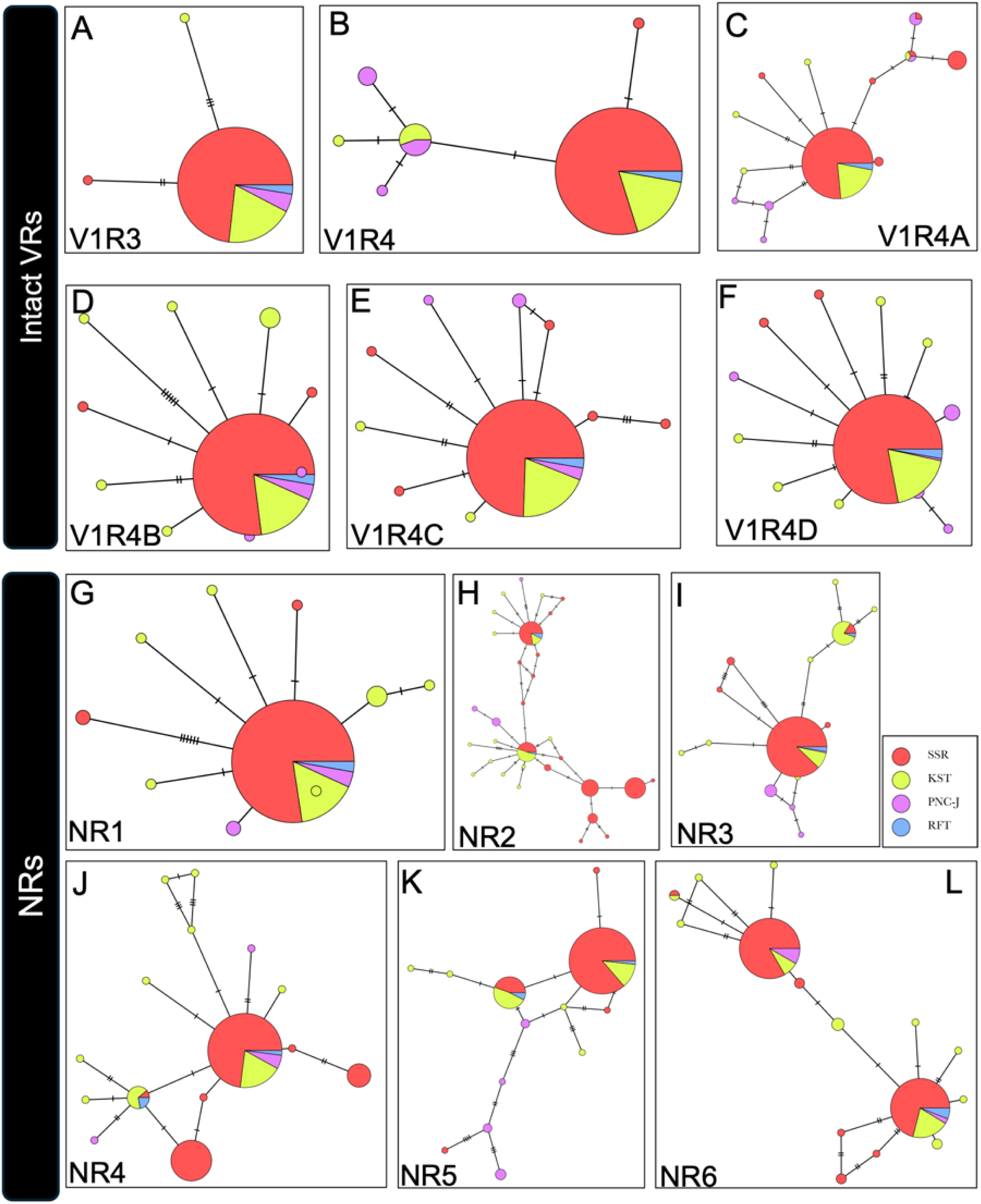
Haplotype minimum spanning networks for six representative intact *VRs* and six representative NRs. Each node indicates one haplotype and each color represents one population. The pie charts within each node show the distribution of the haplotype across the four sampled populations. Each hatchmark in the lines connecting the nodes represents one mutation, such that the length of the connecting lines show mutational distance between haplotypes. A. Haplotype minimum spanning network of intact and disrupted haplotypes of *V1R3* gene. B. Haplotype minimum spanning network of intact and disrupted haplotypes of *V1R4* gene. C. Haplotype minimum spanning network of intact and disrupted haplotypes of *V1R4A* gene. D. Haplotype minimum spanning network of intact and disrupted haplotypes of *V1R4B* gene. E. Haplotype minimum spanning network of intact and disrupted haplotypes of *V1RC* gene. F. Haplotype minimum spanning network of intact and disrupted haplotypes of *V1R4D* gene. G - L. Haplotype minimum spanning networks for six representative NR regions.

### Chemosensory behavioural repertoire and contexts of occurence

Olfactory behaviours with putative relevance to social communication were relatively common in white faced capuchins, occurring an average of 67.68 times/individual/12 hour day (min = 5.82, max = 314.85, sd = 50.61). We recorded 23 distinct behaviours, which we classified into six categories based on whether they were putative signaling or receiving behaviours, and whether they involved environmental substrates, the animals’ own bodies, or the bodies of others: 1) scent marking; 2) urine and substrate investigation; 3) auto body odour investigation; 4) conspecific body odour investigation; 5) anointing with environmental odours; and 6) other. We did not include passive behaviours, such as lying on a branch, even though they might result in the deposition of olfactory signals/cues. A comprehensive ethogram and data on the frequencies of these behaviours, overall (N = 3022.75 contact hours) and within focal follows only (N = 789.95 hours) are provided in Table 2.

### Context and examples of representative behaviours

We observed olfactory behaviours across a variety of social contexts and spatial and temporal scales that are relevant to intra- and intersexual communication. Below we provide detailed examples of many of the behaviours described in Table 2. We organize these according to potential relevance for intrasexual, intersexual, and intergroup communication and also report the presence of seasonality in behaviours. Our goal in doing so is to inspire hypothesis development about evolutionary functions and selective pressures shaping putative olfactory signalling and receiving behaviours so that they can be rigorously tested in future studies. These anecdotes were described by observers to provide examples of extended context for some of the recorded behavioural patterns quantified above.

#### (i) Putative intrasexual communication

On multiple occasions, we observed juvenile and adult males to approach an adult male after he urine washed, then also urine wash in proximity (in contact or within 1-10m) within moments of the initial urine wash. Following some of these instances, the two males engaged in grooming or other affiliative behaviours (e.g., vocalizations, social resting). We have also observed males approach locations where other individuals had urine washed or been resting, then sniff, face-rub, lick or perform the flehmen response to the area for prolonged periods (i.e., 10-30 seconds of inhalation, Supple Video 1). Once, we observed an alpha male urine wash and then present his hand to another adult male to sniff. Another time, an adult male was urine washing as a juvenile male approached and placed his hand below the male to catch his urine. The juvenile then smelled his hand and the adult male mounted him. Other times, a male was observed sniffing or sucking on another male’s anogenital region, which may lead to contact with urine and other odorous compounds.

Adult females, especially kin such as sisters or mothers and daughters, often engage in eye poking and hand sniffing behaviours with each other during which one individual places their fingers in the eyes, nose, or mouth of the other (Supplementary Video 1). This behaviour has been interpreted as a signal of trust between the two individuals (Perry & Smolla, 2020), but it may also have a signalling role. Moreover, adults often sniff or lick infants of the same sex in this species (Perry et al., 2003). We frequently observed individuals sniffing and licking the bodies or urine of infants that were not their own. Additionally, we have observed multiple juvenile males sniffing the bodies (hands, stomach fur) of adult males while appearing to inhale.

Urine washing may also be a behaviour that is more common during times of high excitement or stress (e.g., intergroup encounters, predator encounters, supplants, agonism, sexual interactions). On a few occasions, we observed both males and females urine wash immediately after receiving aggression from other group members (e.g., after being chased or threatened). We have also observed alpha males urine wash repeatedly following predator encounters (e.g., boa constrictors) and/or repeatedly during increased vigilance, and subordinate males urine wash repeatedly while threatening and dropping branches. Finally, we have observed instances of urine washing occurring when one individual supplants another over resources (e.g., fruit trees, dead wood holding insects), most commonly performed by the alpha male (i.e., the supplanter), but also sometimes by the supplanted individual.

#### (ii) Putative intersexual communication

Both adult males and females display olfactory behaviours that appear to be targeted at, or in response to, the other sex. We commonly observed adult males approaching adult females and urine washing within close proximity to them. In one memorable instance, an alpha male rapidly ran ahead of the group (>50 meters) to catch up with an adult female traveling alone and urine washed after approaching to within 5 meters of her, while no other individuals were in view. In other instances we have observed males and females urine wash immediately before or after mating. We often observed males sniffing or licking the urine of females or the locations where they were recently known to be resting (sometimes multiple males will investigate the same urine or location). On multiple occasions after investigation, we observed males resting in the location that they had been intently sniffing. They would then sometimes initiate grooming and licking the female’s anogenital region or leg, or direct a sex display towards her (which includes pacing and gazing at the female while making a “duck face” with pursed lips, often accompanied by sex squeaks or grunts). On more than one occasion, we suspected that the female whose odour was being investigated may have been cycling and fertile because she was not lactating or visibly pregnant and we also observed multiple males following her at close range and directing sex displays toward her exclusively over a period of a few days. Moreover, we have observed females sucking and licking the hands and tail of an alpha male, and sniffing or head-rubbing the urine of other males (the alpha male, a subordinate or a new immigrant male).

#### (iii) Putative searching behaviours and inter-group communication

Olfactory communication can also occur over longer temporal scales, when individuals are not in sight of each other or separated from their group. We commonly observed individuals who were separated from their group sniffing along branches while emitting conspicuous “lost calls”, appearing to search for the direction of their group. On one occasion, when members of his group did not vocally respond to his calls, an isolated male became visibly agitated and he began more rapidly engaging in this sniffing behaviour. In other contexts, during intergroup encounters with neighboring social groups, we commonly observed adult males urine washing and sniffing up and down tree branches, sometimes in pursuit of the other group as they fled, and other times when directly facing off with the other group in a given location. We have also recorded males urine washing with greater frequency after intergroup encounters when they returned to the rest of their group. Finally, on multiple occasions we have observed groups follow the same route that another group was known to have taken the day before, sniffing the branches with increased frequency compared to usual, and often engaging in vigilance behaviours as they travelled along the route. These behaviours are consistent with intergroup olfactory signalling.

#### (iv) Seasonality

Given the reproductive seasonality in this species, and the strong seasonal weather patterns in SSR that might affect signal propagation and reception, we investigated whether seasonality in olfactory behaviours was present in our study population. Season was found to have a significant effect on the daily rates of multiple putative chemosensory behaviours (Figure 5; See Supplementary Table 5 for model results). Urinating, head rubbing on substrates, and sniffing locations occurred more often in the rainy season (Figure 5a-c), while standard urine washing and wind catching occurred more often in the dry season (Figure 5d-e). Vigorous urine washing showed a marginal seasonal effect and occurred more often in the rainy season, though due to few observations, this result should be interpreted cautiously (Supplementary Table 5).

**Figure 4.**
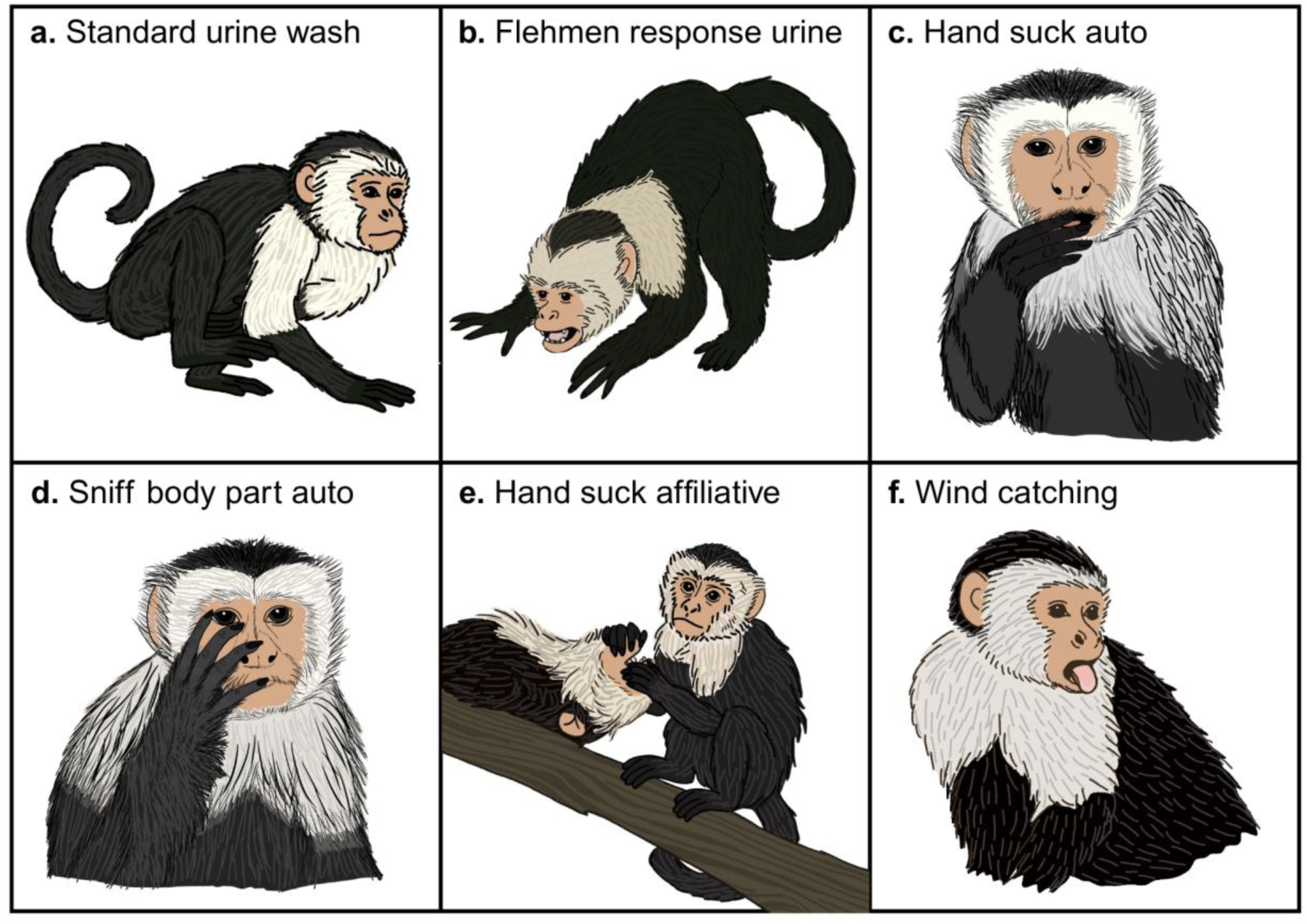
Illustrations of select behaviours observed in wild white-faced capuchins with putative relevance for olfactory social communication. The detailed descriptions of each behaviour are provided in Table 2. a) Standard urine wash; b) Flehmen response urine; c) Hand suck auto; d) Sniff body part auto; e) Hand suck affiliative; f) Wind catching. Artwork by Jordie Hoffman.

**Figure 5.**
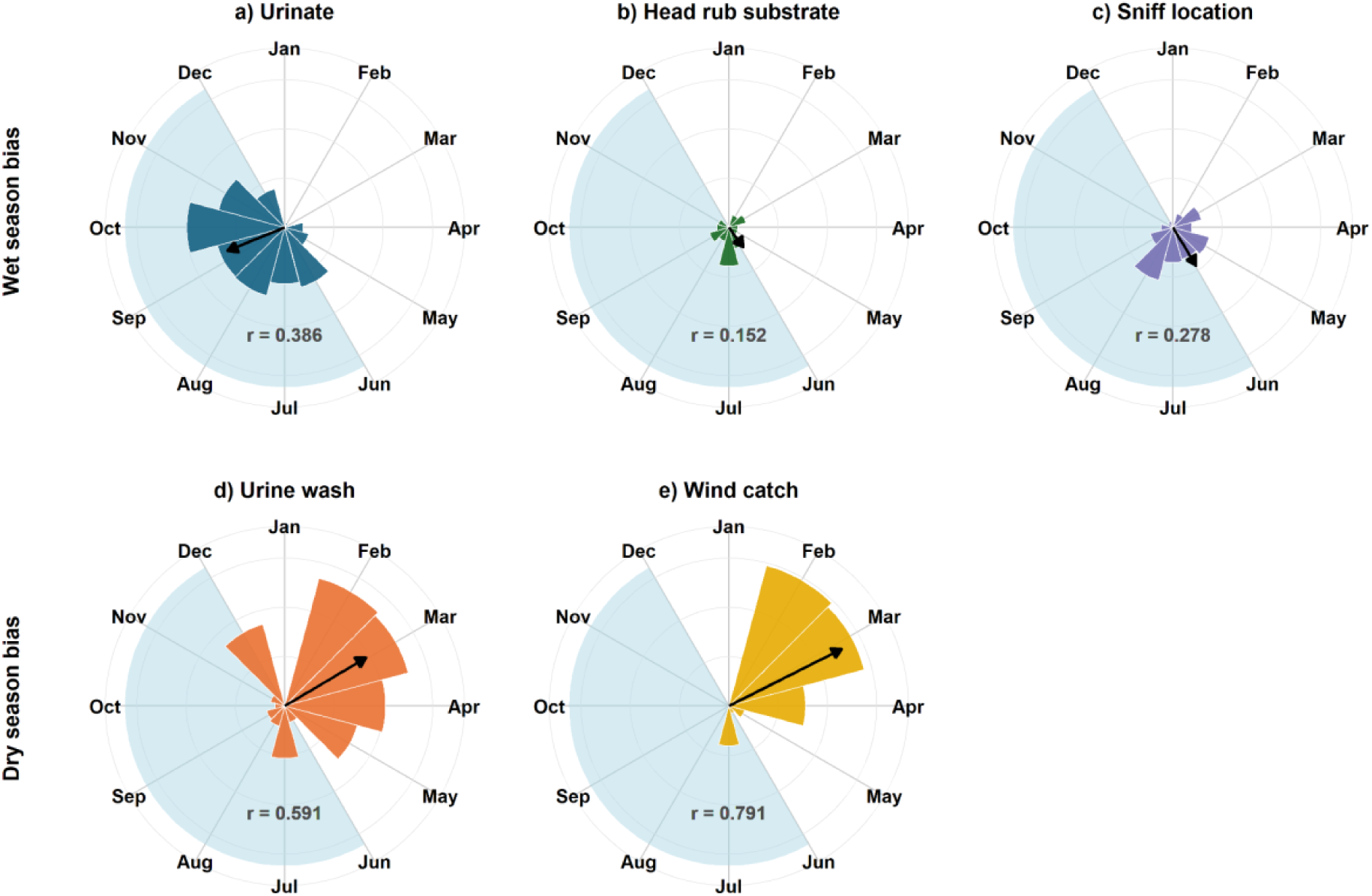
Circular monthly plots for olfactory-associated behaviours with seasonal distributions according to model results. The x-axis represents the day of the year in radians and the y-axis represents mean daily rates (count/12 hour day) of olfactory-associated behaviours observed within focal follows (789.95 focal hours).The blue shading within the histograms represents the duration of a typical 6-month rainy season in Sector Santa Rosa, Costa Rica during our 2021-2024 study period, characterized as cumulative rainfall >100 mm in the 30 days prior to each sampling date (Hogan & Melin, 2018), and extending from around June-December. In each histogram, the orientation of the mean vector arrow represents peak seasonal timing and its length (r; 0-1) indicates the strength of behavioural seasonal concentration, with longer vectors representing greater seasonality.

## Discussion

Our central aim was to assess the potential role of social olfaction in white-faced capuchins. We first sought evidence for the presence of putative functional *VR* genes using sequencing data from monkeys living in four geographically distinct locations and then investigated whether patterns of genetic diversity in these *VR* genes are consistent with selective constraints. Secondly, we assessed behavioural data for evidence that olfaction is used regularly in socially-relevant contexts in this species. Overall, we found evidence consistent with evolutionary constraint on intact vomeronasal receptor genes and documented many behaviours with chemosensory communication potential. These results support a potential role for olfaction in social communication in a species of wild platyrrhine primate. In particular, they indicate that there are many opportunities for chemosensory signalling in diverse contexts, highlighting a role of odours and olfaction in intra- and intersexual communication, competition, and reproduction.

### Aim 1. VR genes and signatures of evolutionary constraint

Our study provides the first population-level investigation of the *VR* gene repertoire and evolutionary dynamics in wild monkeys. We provide multiple, complementary lines of evidence that intact *VR* genes in white-faced capuchin monkeys are evolving non-neutrally. We demonstrate that at least 12 *VR* genes are intact and exhibit signatures consistent with evolutionary constraint. We further show that disrupted haplotypes were rare for the intact *VR* genes we identified in white-faced capuchins, and both dominance of a single intact haplotype and the reduced Tajima’s D values compared to NRs are consistent with evolutionary constraint. If so, selection may be acting on these genes to maintain receptor integrity. This pattern aligns with broader evolutionary trends in anthropoid primates, where reductions in sensory gene repertoires are often accompanied by the retention of a small number of functionally important loci under selection (Hiwatashi et al., 2010; Niimura et al., 2018; Wooding, 2011). Non-neutral evolution have been inferred for other chemosensory and sensory receptor genes in primates (Akhtar et al., 2022; Hiwatashi et al., 2010; Wooding, 2011), including olfactory receptors, taste receptors, and opsin genes, highlighting a broader pattern in which sensory systems are shaped by selective pressures despite reduced gene repertoires (Niimura, 2012; Niimura et al., 2018; Niimura & Nei, 2006).

The functional status of *VRs* in platyrrhines is of ongoing interest, with some studies suggesting extensive degeneration of the vomeronasal system in anthropoid primates (i.e., the major primate clade that includes platyrrhines: Niimura, 2012; Niimura & Nei, 2006; Yoder & Larsen, 2014), including its loss in many catarrhines (“Old World monkeys and apes), yet persistence of a vomeronasal organ (VNO) and multiple *VR*s in platyrrhines (Moriya-Ito et al., 2018; Smith et al., 2011). Variation of VNO size, location and of *VR* repertoires in platyrrhines may be extensive but is poorly understood (Smith et al., 2011). For example, *VR* repertoire has been investigated in only two other platyrrhine species (Moriya-Ito et al., 2018). Common marmosets, a small platyrrhine species that frequently makes and investigates scent marks (Carboni et al., 2026), have 7 intact *VR*s, and black-capped squirrel monkeys have as many as 15 (Moriya-Ito et al., 2018, Yoder & Larsen, 2014; Young et al., 2010). All intact *VR* genes we identified in white-faced capuchins belong to the *V1R* gene family; we did not find any intact *V2R* genes, which is consistent with the other studies of platyrrhine species. Overall, our results here suggest that capuchins may rely on a small number of functionally important, evolutionarily constrained receptors similar with other platyrrhines, rather than maintaining a large and rapidly evolving VR repertoire, as seen in strepsirrhines (Niimura et al., 2018; Smith et al., 2007).

The number of intact *VR* genes identified in capuchins is more similar to the *VR* repertoires reported for other platyrrhine primates. Though there are only a few reports on *VR* repertoire among platyrrhines, there have been many investigations on *VR* repertoire in catarrhines and all of them show a relatively very small number of intact *VRs*. Among catarrhines studied to date, one intact *VR* gene has been identified in golden snub-nosed monkeys (*Rhinopithecus roxellana*) and green monkeys (*Chlorocebus sabaeus*); two intact *VR*s have been identified in bonobos (*Pan paniscus*), northern white-cheeked gibbons (*Nomascus leucogenys*) and western gorillas (*Gorilla gorilla*); four intact *VR* genes have been annotated for the common chimpanzees (*Pan troglodytes*); and five have been identified for Sumatran orangutans (*Pongo abelii*) (Moriya-Ito et al., 2018; Young et al., 2010). Based on recent genomic analyses, crab-eating macaque (*Macaca fascicularis*) and rhesus macaque (*Macaca mulatta*) genomes appear to carry no intact *VR* genes (Moriya-Ito et al., 2018). Given that capuchins are reported to have a VNO, which is thought to be broadly absent catarrhine monkeys (Smith et al., 2011). In addition, the number of reported *VR* pseudogenes varies several fold among primate species, painting a complicated evolutionary picture. The picture is further complicated by the expression patterns of *VR* genes, which can be found both within and outside of the VNO in platyrrhines and perhaps other mammals (Hills et al., 2024; Moriya-Ito et al., 2018; Yakubets et al., 2025). Future transdisciplinary study of behaviour, anatomy, and molecular biology of primate *VR*s will hopefully shed new light on their putative roles in olfaction and beyond.

The persistence of intact *VRs* under evolutionary constraint suggests that, despite the overall reduction of the vomeronasal system in anthropoid primates relative to other mammals (Yoder & Larsen, 2014), these receptors likely retain biologically relevant functions in capuchins. The specific function of distinct capuchin *V1R* genes remains to be elucidated. Nevertheless, given the extensive social functions in other mammals, detection of pheromones or other social odours may be likely. Living in gregarious female philopatric multimale-multifemale groups, capuchins lead complex social lives wherein individuals preferentially affiliate with kin (Perry, 1996a; Schoof & Jack, 2014), select mates (Buckner et al., 2021), and navigate the social complexities that come with natal dispersal, emigration, and male-male competition over the sole alpha male position in each group. Each of these tasks could plausibly involve chemical cues. Furthermore, behavioural and genetic evidence support inbreeding avoidance in wild capuchins (Godoy et al., 2016), and olfaction might play a role here as well, as it does in other mammals (Boulet et al., 2009; Henkel & Setchell, 2018; Mitchell et al., 2018). The dominance of a single *VR* haplotype, for intact genes, across ecologically distinct populations suggests that these receptors may be involved in core social or physiological processes rather than fine-scale environmental adaptation. Identifying the specific ligands for capuchin *VR*s, and comparative analyses across diverse platyrrhine species (especially callitrichids who heavily engage in scent marking and other chemosignalling behaviours: Epple et al., 1986; Heymann, 2022; Snowdon & Ziegler, 2021), and other mammals, would shed additional light into the evolution and adaptation of *VRs* and social olfactory communication in monkeys.

### Aim 2. Behaviours and putative olfactory communication

During our field study, we observed and characterized an extensive repertoire of putative chemosensory behaviours performed by white-faced capuchins. Many of these behaviours were performed frequently (> 5 times per day), closely mirrored those used for communication by other more macrosmatic mammals (Brown & Macdonald, 1985; Johnson, 1973; Wilson, 1971; Wyatt, 2014), and followed seasonal patterns, collectively suggesting a role in responding to spatiotemporal social and/or ecological challenges. Capuchins are moderately seasonal breeders (Brasington et al., 2022), so it is possible that some behaviours, such as urine washing and the olfactory investigation of urine and substrates, may be involved in reproductive communication. While behaviours such as wind catching and urine washing have been suggested to aid in seasonal thermoregulation (Campos & Fedigan, 2009; Schmidt & Seitz, 1967; but see Campos et al., 2007) and may therefore track seasonal increases in water consumption, they may nonetheless be socially informative. The dry season is also much windier than the rainy season, which might influence the efficacy of “wind catch” behaviours for thermoregulation, communication, or potentially other motivations. Similarly, sucking body parts and anointing fur with highly fragrant plants might conceivably play a primary role in removing or repelling ectoparasites (Baker, 1996; DeJoseph et al., 2002; Freeland, 1981; Hart & Hart, 2018), while also presenting informative olfactory cues involved in social bonding or other communication (Leca et al., 2007; Supplemental video S1). Integrating these behavioural observations with confirmation of evolutionary constraint in *VRs* provides powerful evidence of the functional importance of olfactory communication in this species.

Urine washing was one of the most commonly recorded behaviours in SSR during our study. A number of functional explanations have been proposed for urine washing (e.g. thermoregulation, the improvement of grip to substrates, and resource marking around fruit trees; Campos & Fedigan, 2013). In line with other studies, we found that standard urine washing in male capuchins followed seasonal patterns and was performed most often in the dry season (Campos et al., 2007; Campos & Fedigan, 2013; Poirier et al., 2025), when most male emigrations and alpha male replacements also occur (Brasington et al., 2022; Fedigan, 2003). Given that urine washing frequency varies with sex and male dominance rank status (Campos et al., 2007; Campos & Fedigan, 2013; Poirier et al., 2025) and is performed and investigated by both sexes, it may also be involved in intrasexual and intersexual communication, a possibility supported by the observations reported here. If urinary odours are informative with respect to dominance rank, health, and/or other markers of competitive ability, chemosignals may influence mating and male dispersal behaviour. Additionally, female capuchins have visually cryptic ovulation (Fedigan & Jack, 2013) and most conceptions in SSR occur during the rainy season (Brasington et al., 2022), when we observed the highest rates of location sniffing by males. If female urinary odours are informative with respect to female reproductive status, this may, in turn, influence male reproductive strategies (Smith et al., 1997; Smith & Abbott, 1998; Washabaugh & Snowdon, 1998). Interestingly, urine washing is noted to be less common at other white-faced capuchin study sites (e.g., BCI, Jicaron; pers comms MC and BB), suggesting environmental or social drivers of variation that may be interesting to pursue in future research.

The seasonal use of olfactory behaviours may also be related to climatic variables. For example, while odour detection increases in humid conditions (Kuehn et al., 2008; Sen Majumder & Bhadra, 2015), rainy conditions may limit signal persistence by diluting odorants directly upon contact (Alberts, 1992) and/or indirectly by diluting urine via increased water consumption. If so, capuchins may need to get closer to substrates in order to detect odours, contributing to the higher rates of sniffing observed in the rainy season. In contrast, urinary odours may last longer on substrates during the dry season due to the lack of rain, which speeds degradation (Alberts, 1992). Groups also spend prolonged periods of time resting in shaded trees and in close proximity to rare and coveted water sources when it is drier (Campos & Fedigan, 2009), potentially facilitating olfaction as a medium for communication. While we did not explicitly find any urinary investigation behaviours to be seasonally distributed, this may be related to the rarity of observing paired sequences of urination followed by investigation from another individual.

### The importance of social olfaction to platyrrhines

Historically, many inferences on the relevance and use of mammalian olfactory communication have been based on morphological characteristics (Doty, 2003; Eisenberg & Kleiman, 1972; Meisami & Bhatnagar, 1998; Smith & Rossie, 2006). Platyrrhines lack some of the morphology traditionally used to interpret the relevance and function of mammalian olfactory communication, such as large olfactory bulbs and a wet rhinarium to aid in trapping and dissolving odors for enhanced processing by olfactory receptors and *VRs* (Niimura et al., 2018; Smith et al., 2007). Yet, our findings provide clear evidence of a persistent role of olfaction in capuchin social lives. By adding data on *VR* genes and chemosensory behaviours we complement previous studies of olfactory morphology to broaden the understanding of the platyrrhine olfactory landscape. Our data challenge the historical notion that anthropoids minimally rely on olfactory communication to navigate their social environments (Gilad et al., 2004; Heymann, 2006b). Indeed, while the specialized scent glands and glandular scent marking behaviours characteristic of callitrichid platyrrhines (Epple et al., 1986; Heymann, 2006a; Snowdon & Ziegler, 2021) are not ubiquitous across the platyrrhines, species of other platyrrhine families, such as the atelids, retain scent glands that may produce odours involved in social signalling (Epple & Lorenz, 1967; Salazar et al., 2015). Furthermore, platyrrhines seem to possess a vomeronasal organ (some with derived morphology; Smith et al., 2011) and many engage in olfactory investigation and urine washing as a putative chemosignalling behaviour (e.g., woolly monkeys, spider monkeys, owl monkeys, and howler monkeys; Heymann, 2006a).

Our research adds to evidence from lemurs and other strepsirrhines that olfactory behaviours in primates, such as sniffing and scent marking, may be linked to *OR* and *VR* repertoires (fat-tailed dwarf lemur: Yoder et al., 2014; mouse lemurs: Aujard, 1997; northern greater galago: Moriya-Ito et al., 2018). By examining behaviour and evidence of *VR* functionality in tandem, our results can be carried forward to substantiate inferences on olfactory capability based on vomeronasal complex morphology and behavioural observation, filling a much needed gap in piecing together the evolutionary history of these traits in primates. Studies of intact VR repertoires and olfactory-associated behaviours in platyrrhines are exceedingly rare. To date, the *VR*s of only two other platyrrhines have been studied (Moriya-Ito et al., 2018) but population data are sparse. The expansion of these datasets across platyrrhines and other primates holds great promise in illuminating not only selective pressures driving trait preservation in primates, but also those driving analogous reductions in olfactory gene repertoires across mammals (bats: Zhao et al., 2011; cetaceans: Yu et al., 2010; primates: Liman & Innan, 2003) and contributing to our overall understanding of sensory landscape evolution.

Communication in animals is inherently multimodal, though with potential tradeoffs between increased reliance on vision and olfaction (and other sensory modalities) across mammals (Chi et al., 2025; Higham & Hebets, 2013; Kawamura & Melin, 2017). The frequency with which diurnal, group-living monkeys use olfaction is perhaps surprising and suggestive of a persistent role of olfaction maintained by natural selection as part of a suite of sensory modalities used for communication. Capuchins have acute vision and polymorphic trichromacy (Hiramatsu et al., 2017; Kawamura & Melin, 2017) and complex vocal repertoires (Gros-Louis et al., 2008), motivating an open question about how olfaction fits into in their multimodal communication repertoire. Olfactory signals persist over relatively long periods of time (Finnerty et al., 2022; Uy & Safran, 2013; Williams et al., 2023) and may be particularly important for platyrrhine primates that, as a whole, are more arboreal than their cercopithecine cousins and often have their lines of sight obscured by foliage. Olfactory signals may additionally enhance or modulate signal efficacy when paired with acoustic and/or visual cues and signals (Partan & Marler, 1999). Behavioural observations provide a potential window into putative functions.

Based on the social contexts of our field observations, we predict that olfactory communication in capuchins, as in other mammals, may play a role in signalling and reinforcing dominance ranks (Borak et al., 2025; Cafazzo et al., 2012; Hurst et al., 2001; Poirier, 2019; Setchell et al., 2010; Walker-Bolton & Parga, 2017), facilitating reproduction in a species with cryptic female cycling (Baum & Cherry, 2015; Beauchamp, 1973; Chang et al., 2000; Kean et al., 2011; Poirier et al., 2021; Powers et al., 1979), communicating individual, group and species identity (Carboni et al., 2025; Henkel et al., 2015; Henkel & Setchell, 2018; Poirier et al., 2021; Setchell et al., 2010; Vaglio et al., 2016), and possibly in territory and resource defense (Biswas et al., 2024; Black-Decima & Santana, 2011; Garcia de la Chica et al., 2024; Jordan et al., 2010; Miaretsoa et al., 2022; Remonti et al., 2011).

### Conclusion and Future Directions

This study adds to our understanding of *VR* repertoires and patterns of selection in wild monkeys. Our findings that *VR* genes show evolutionary constraint signatures across multiple populations of white-faced capuchins point to an evolutionarily conserved role of *VR*s in olfactory signalling. Furthermore, that chemosensory behaviours occur at high rates and across many socially-relevant contexts suggests that olfaction plays an active role in capuchin intra- and intersexual communication, competition, and reproduction.

Despite providing insights, our study had several limitations. All genetic inferences in this study relied on short-read whole-genome sequencing data, which can complicate accurate variant calling and phasing in *VR* genes due to sequence similarity among paralogs. Although stringent quality filters were applied and analyses were restricted to high-confidence variants, some rare or structurally complex variants might remain undetected due to methodological challenges. Future studies with long-read sequencing can overcome this challenge. Additionally, gene expression studies of VNO and other tissues to identify where and to what extent *VR*s are expressed would be highly instructive. To build our behavioural ethogram, we combined focal follows and ad lib observations of males. A more standardized and comprehensive study of all age-sex classes, drawing on a consistent set of observation approaches, would therefore further enrich our understanding of inter-individual variation in olfactory behaviours. Finally, the functional consequences of *VR* variation were not experimentally validated. To examine whether capuchin VR genes are involved in social communication, future studies could conduct olfaction experiments with captive individuals, or expose capuchin-sourced volatile and semi-volatile compounds (for example those found in urine) to *VR* proteins expressed *in vitro* within a heterologous cellular expression system (e.g., Baldwin et al., 2014). Overall, we hope our findings will inspire future research and help draw attention to the presence of chemosensory behaviours in capuchins and other primates whose use of olfaction may be underappreciated in social contexts.

## Acknowledgements

We thank the Costa Rican Park Service – including the Ministerio de Ambiente y Energía (MINAE), el Sistema Nacional de Áreas de Conservación (SINAC), and the Comisión Nacional para la Gestión de la Biodiversidad. We give special thanks to Roger Blanco Segura and María Marta Chavarria, for supporting our ongoing research in the Área de Conservación Guanacaste. We thank Saúl Cheves Hernandez for his assistance with project management in Costa Rica, Noah Snyder-Mackler for help with sample processing, and Anja Widdig for feedback on the manuscript and many useful conversations. We thank dedicated field researchers, including Marcos Alvarez-Garita, Nicholas Chapoy, Alice Galotti, Rafael Jiménez Montero, Clara Martinez Suarez, Daniela Salazar Sánchez, Robinson Sandoval Benavides, Giulia Severino, Yarenis Villalobos Abella, and Jonathan Wilkins for their contributions to the capuchin project. We are grateful to Anja Widdig for ongoing discussions on the topic. We also thank MiAmbiente in Panama for permission to conduct research in Panama and the staff at the Smithsonian Tropical Research Institute, and particularly Lucia Tórrez-Herrera for logistical support. Thanks to Pedro Castillo-Caballero, James T. Chaves, and Meredith Carlson for help with sample collection in Panama.

## Ethics statement

This research adhered to the laws of Costa Rica, the United States and Canada and complied with protocols approved by SINAC and the Área de Conservación Guanacaste and by the Canada Research Council for Animal Care through the University of Calgary’s Life and Environmental Care Committee (ACC protocol AC15-0161). Samples were collected with permission from the Área de Conservación Guanacaste (ACG-PI-033-2016).

Sample collection at Taboga was conducted under permit CPl-SlNAC-PNl-ACAT-011-2021. Research at SSR was conducted under permits R-042-2023-OT-CONAGEBIO; R-013-2022-OT-CONAGEBIO; R-019-2020-OT-CONAGEBIO; R-029-2017-OT-CONAGEBIO and R-025-2014-OT-CONAGEBIO. Research at KSTR was conducted under permit numbers R-047-2024-OT-CONAGEBIO and R-002-2020-OT-CONAGEBIO, and Canada import permit numbers CFIA: A-2016-03992-4; A-2018-06070-4; A-2020-00192-4; A-2021-01188-4; A-2022-05488-4; A-2023-06194-1. Sample collected at PNC-J, Panama was conducted under permission from MiAmbiente in Panama (permit # SE/A-6-2020 and ARB-158-2022) and Canada import permit number, CFIA: A-2022-05395-1.

## Data availability statement

The raw sequencing data of study individuals can be accessed at NCBI sequence read archive (SRA) under BioProject ID PRJNA1508398. The variant calls in VCF format are available at Zenodo and can be accessed under DOI no. 10.5281/zenodo.22004309. The variant calls are also available at DRYAD under DOI no. 10.5061/dryad.b2rbnzsx9.

## SUPPLEMENTARY TABLES

S Table 1: List and characteristics of white-faced capuchin study individuals.

S Table 2: A summary of vomeronasal receptor repertoire in white faced capuchin genome.

S table 3: List of identified neutral references in capuchin genome.

S table 4: Depth of each intact, disrupted and truncated VR gene, and NR region in studied capuchin monkeys.

S Table 5: Estimates from a GLMM and LRT assessing the effect of categorical season (rainy/dry) on daily behavioural rates (count/hour). The GLMM reference level is the rainy season. Significant p-values are bolded and season of greatest occurrence is noted, for behaviors where significant seasonality is detected.

## SUPPLEMENTARY Figures

### Legends

**S Figure 1:**
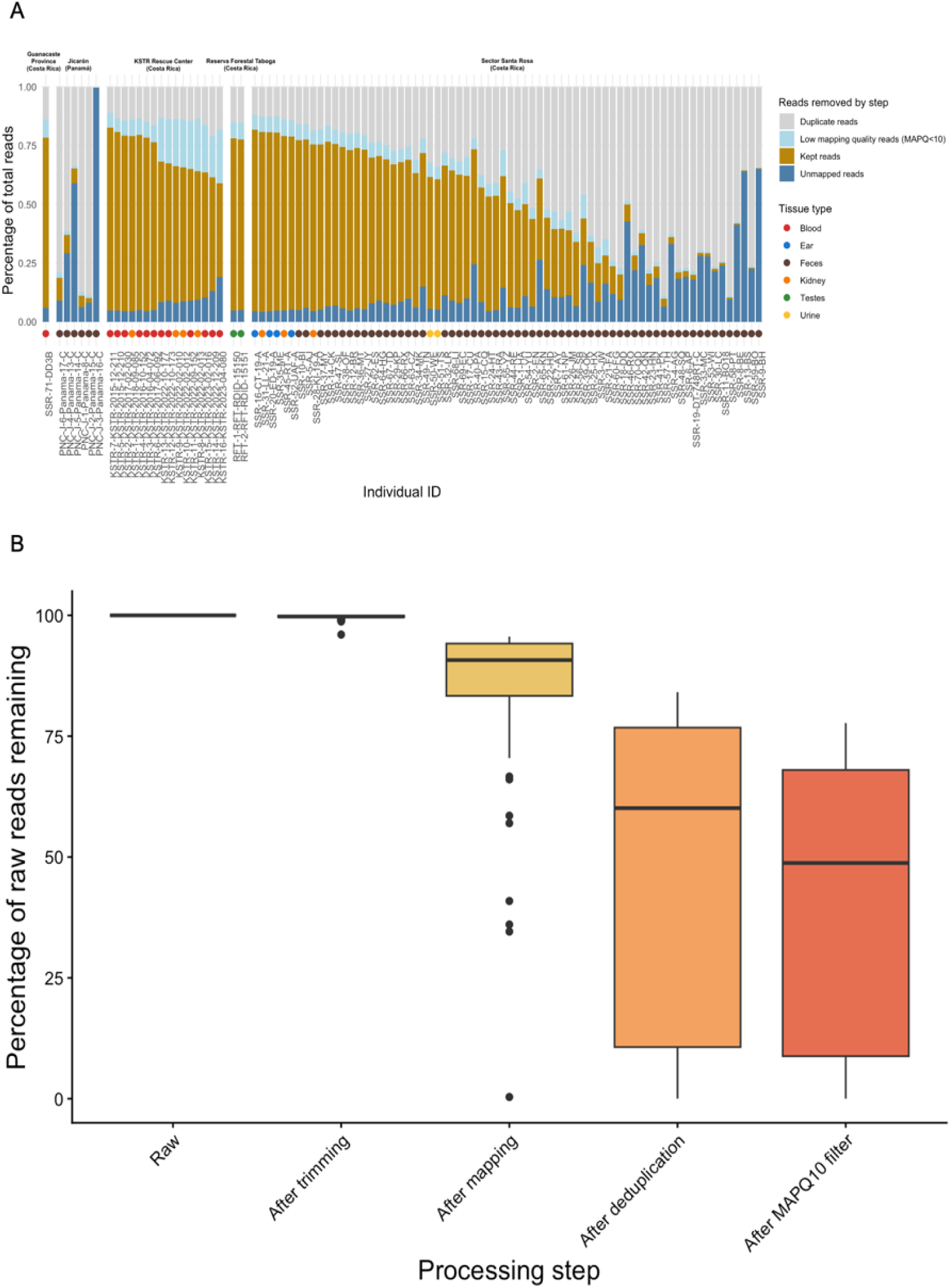
Summary statistics for read quality control, and removal of human contamination. A. Each bar represents one individual included in this paper. The total bar height represents 100% of the reads, and each color indicates the percentage of reads either kept or removed at each processing step. Individuals are grouped by sampling location and color-coded by tissue type. B. Percentage of raw reads kept at each filtering step starting from raw reads to trimming, mapping, deduplication and post-mapping filtration.

**S Figure 2:**
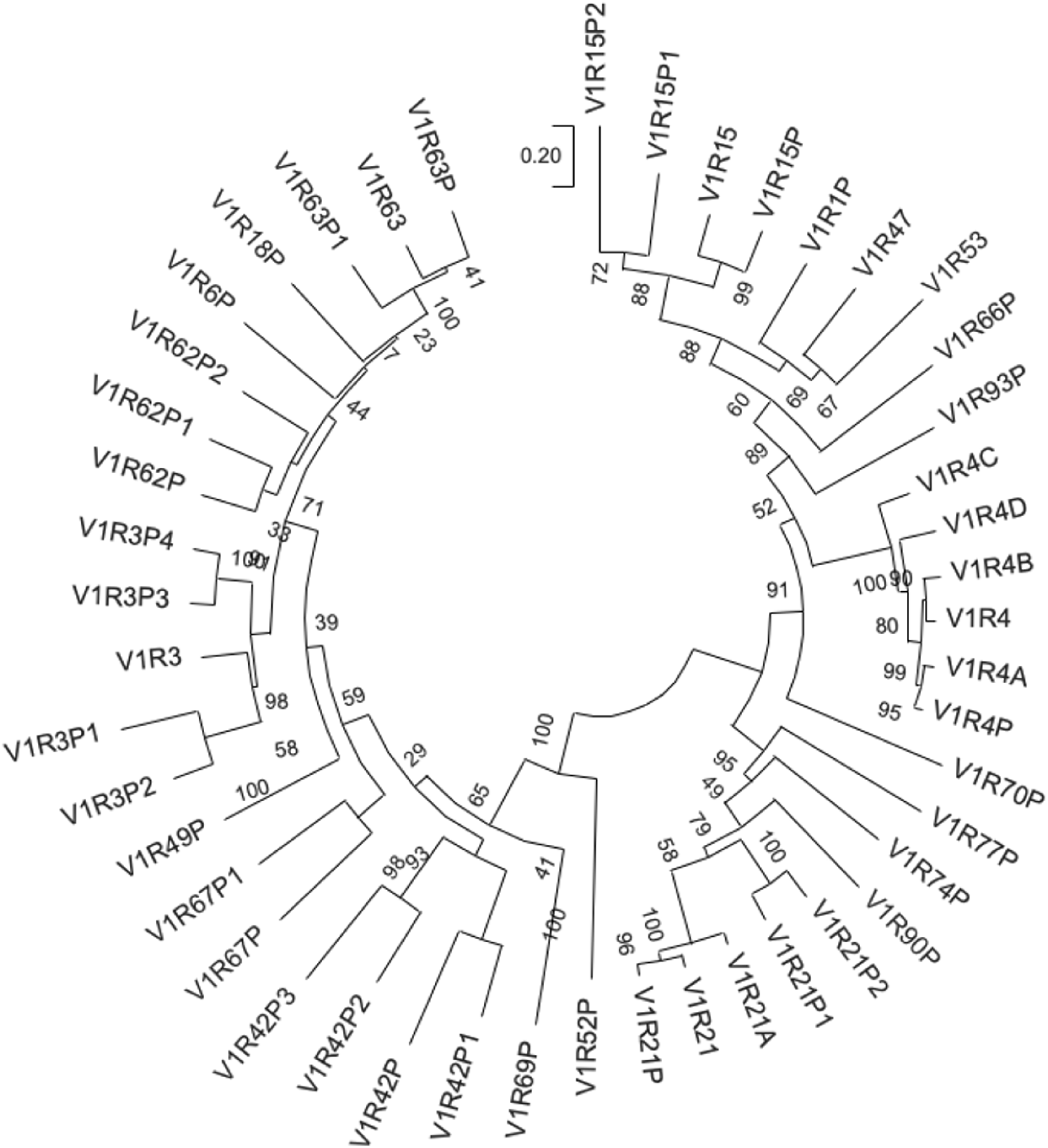
Phylogenetic relationship of *V1Rs*. A map of paralogous *VR* genes within capuchin genome assembly. Any gene with the suffix P is a disrupted gene based on genomic assessment.

**S Figure 3:**
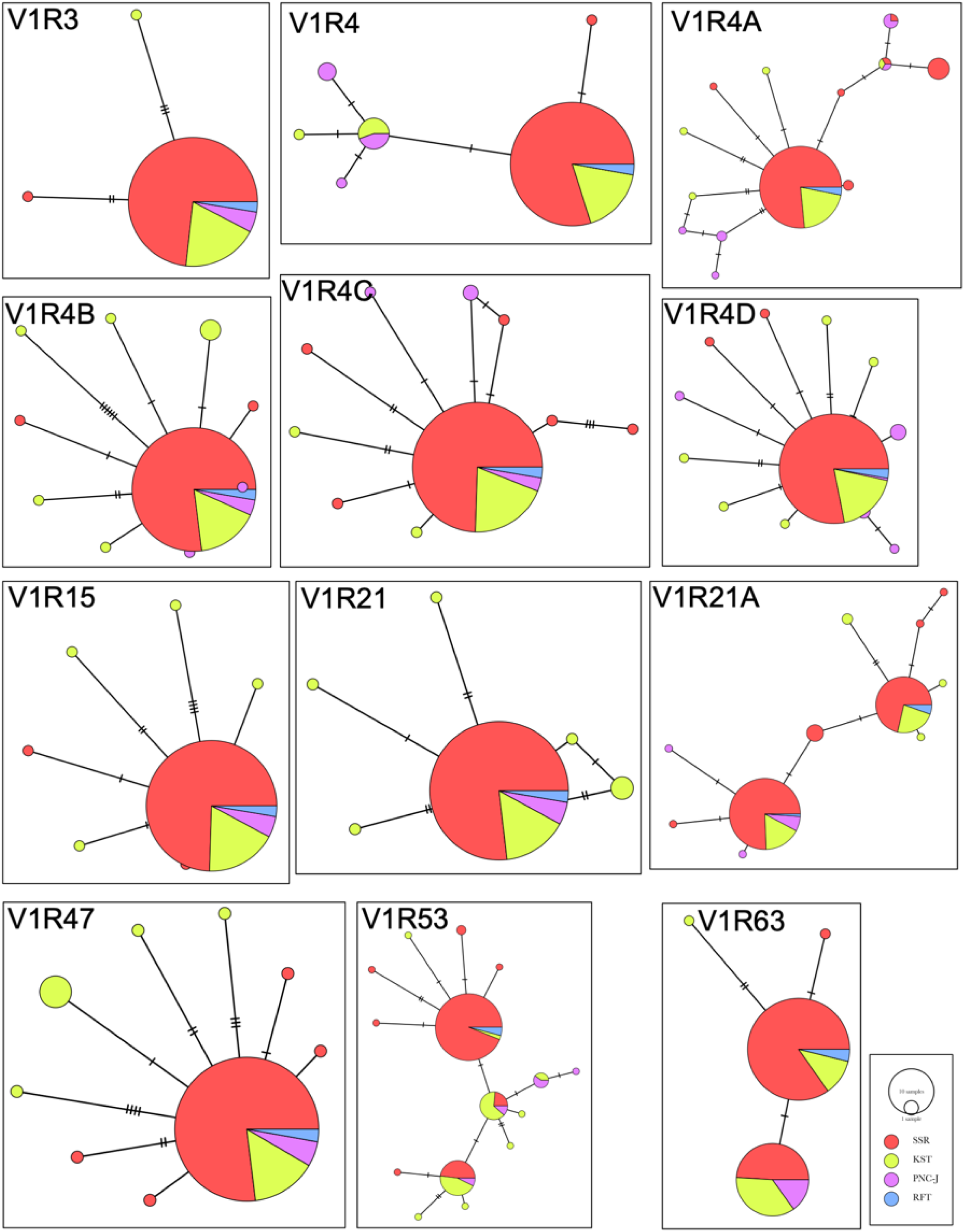
Haplotype minimum spanning networks for 12 intact *VRs*.Each node indicates one haplotype and each color represents one population. The pie charts within each node show the distribution of the haplotype across the four sampled populations. Each hatchmark in the lines connecting the nodes represents one mutation, such that the length of the connecting lines show mutational distance between haplotypes. Haplotype minimum spanning networks for 12 *VR* genes.

**S Figure 4:**
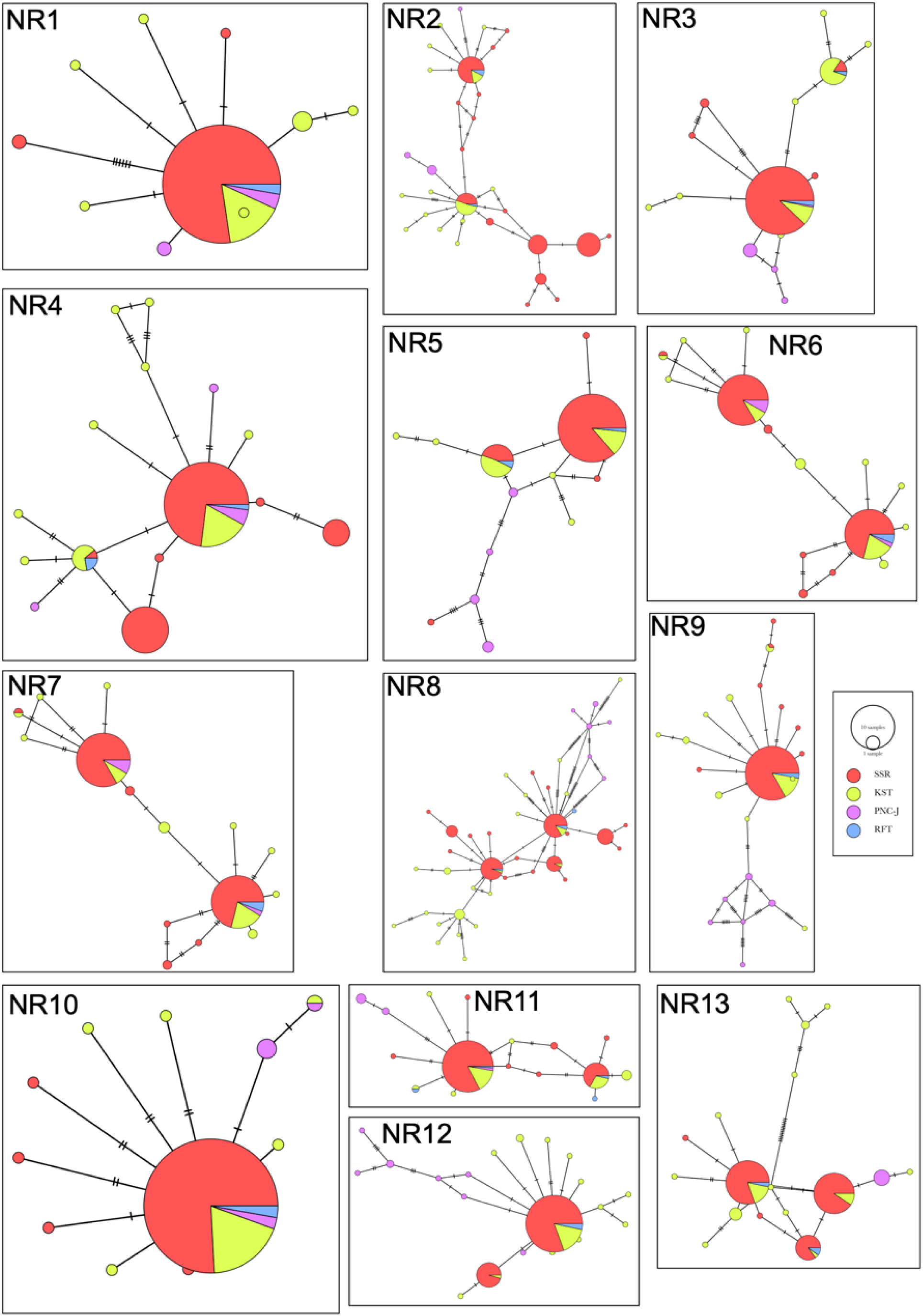
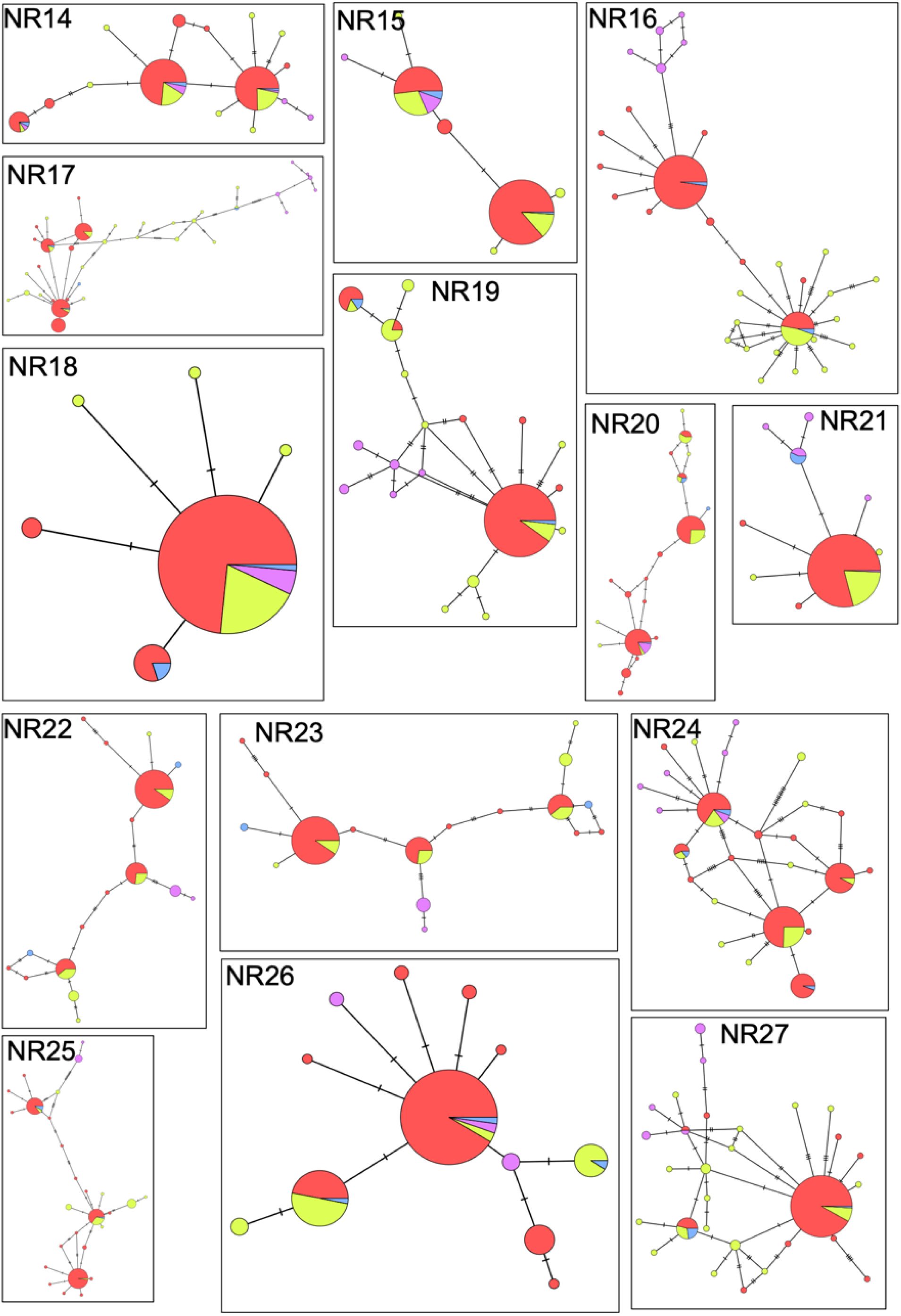
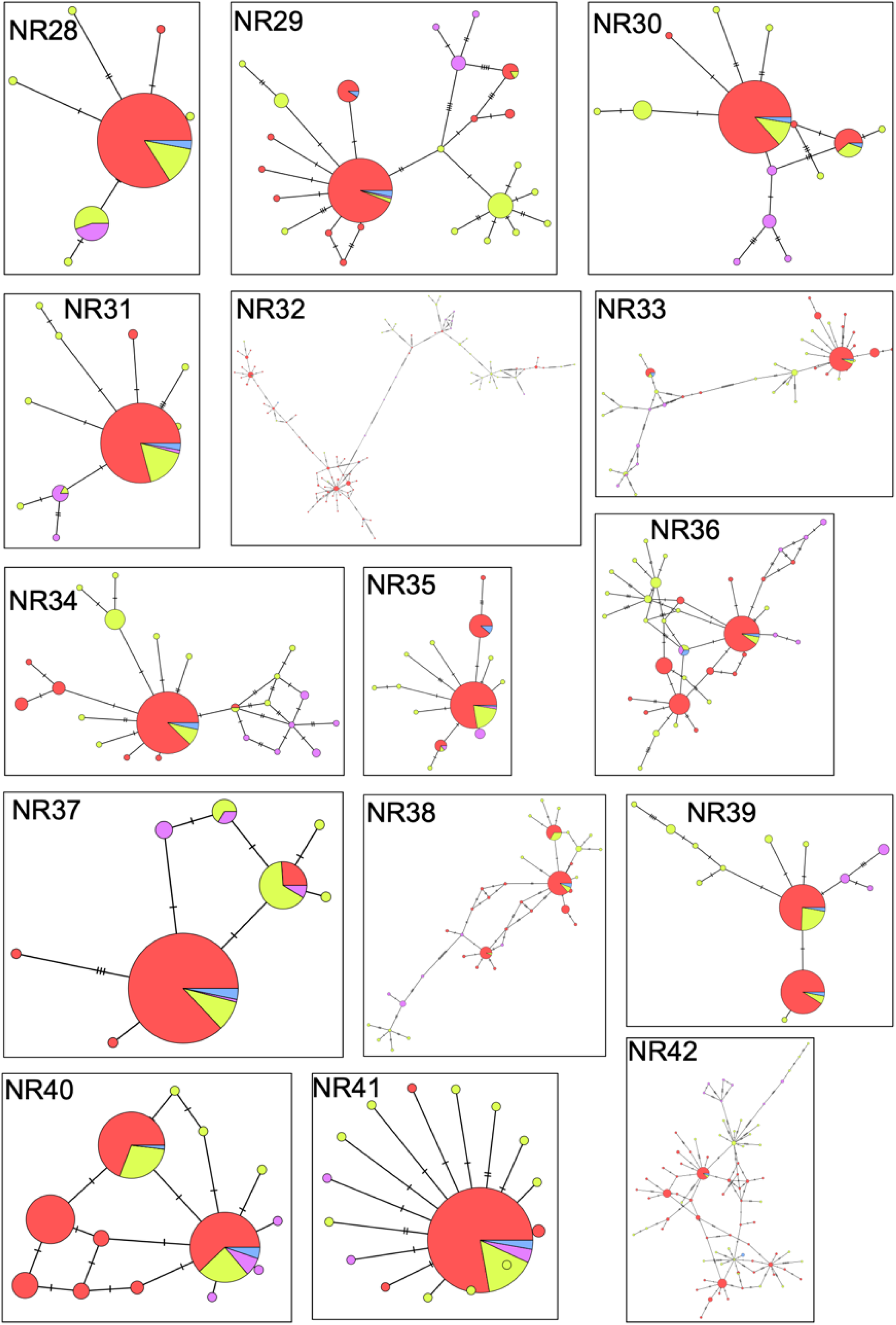
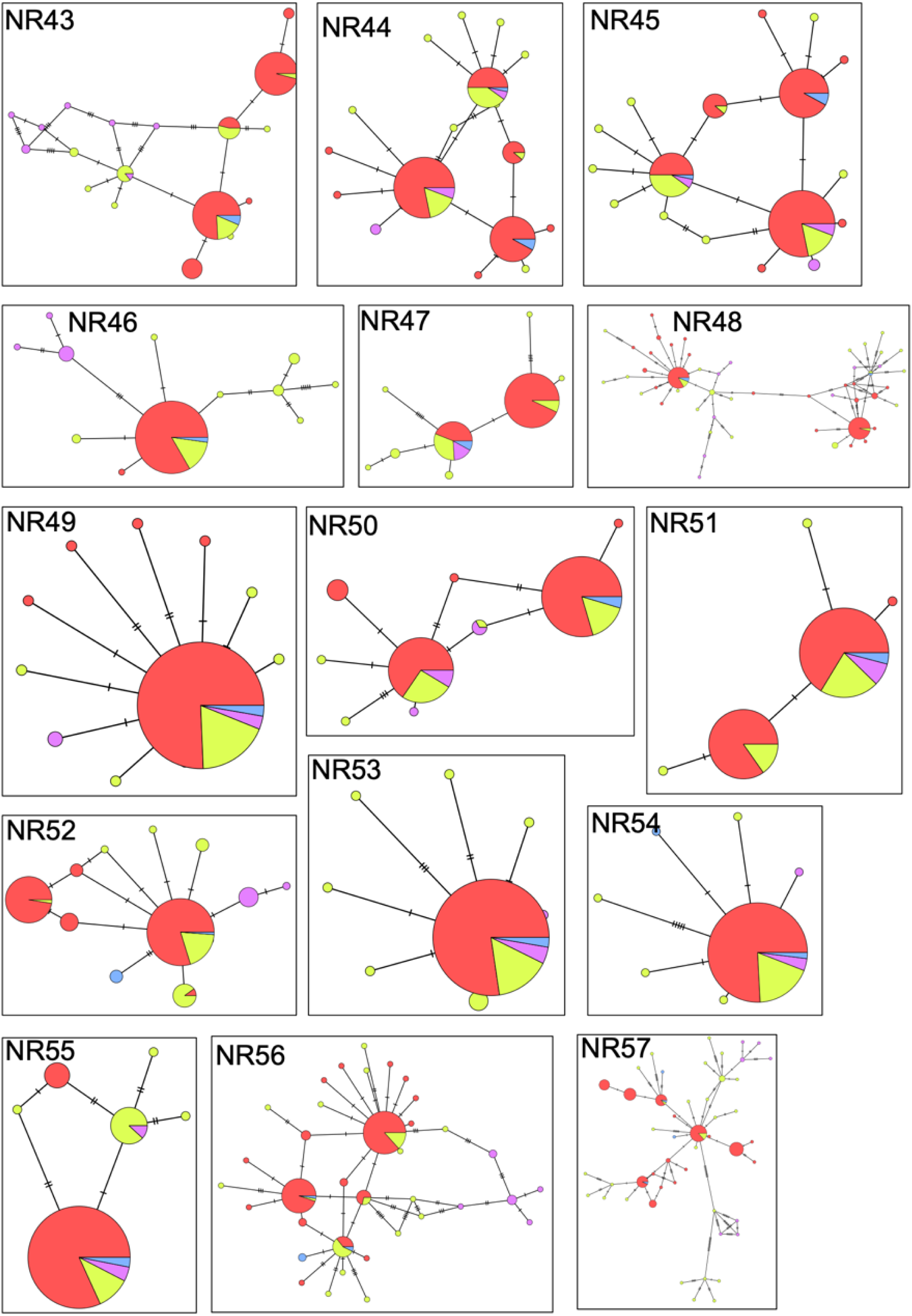
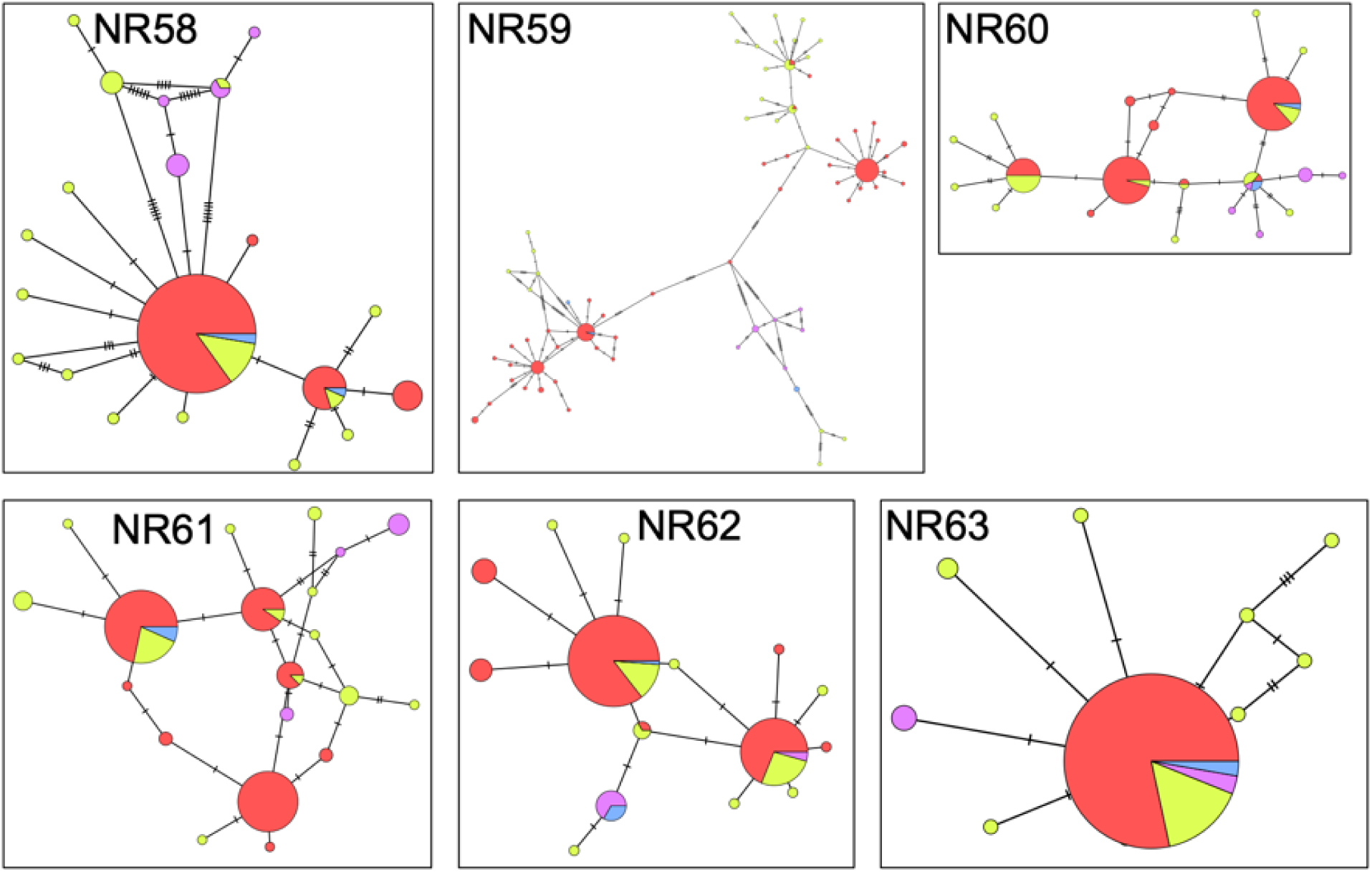
Haplotype minimum spanning networks for 64 NRs, not shown in main text. Each node indicates one haplotype and each color represents one population. The pie charts within each node show the distribution of the haplotype across the four sampled populations. Each hatchmark in the lines connecting the nodes represents one mutation, such that the length of the connecting lines show mutational distance between haplotypes. Haplotype minimum spanning networks for 64 NR regions.

