## Supplementary material for "Genetic and behavioural repertoires support olfactory communication in wild capuchin monkeys": Supplementary Methods.pdf

### 1. Bioinformatics Processing of Reads

First, we used *TrimGalore 0.6.10* (Krueger, 2015) to remove adapter sequences and trim low-quality bases from the ends of all sequencing reads. To assess potential human contamination, we mapped the trimmed reads against the human genome (hg38) and *Cebus imitator 1.0* genome using BBSplit (Bushnell, 2014). Default parameters were used to bin reads that mapped uniquely to the human genome, uniquely to the capuchin genome, or remained unmapped. Reads that mapped equally well to both genomes (ambiguous reads) were assigned to both genomic bins. The proportion of human contamination was calculated as the fraction of reads that mapped unambiguously to the human genome. This contamination check was performed for 75 biological samples; for the subsequent 27 samples, it was not repeated, as initial results indicated minimal human contamination (S Figure 1). Reads assigned to the *Cebus* bin were subsequently mapped conventionally to the capuchin genome.

The reads were mapped to the *Cebus imitator* 1.0 genome assembly with Bowtie2 (Langmead & Salzberg, 2012) and the resulting alignments were sorted using samtools 1.19.2 sort (Li et al., 2009). For samples where competitive mapping against the human genome was skipped, the trimmed reads were mapped directly in the same manner. We removed PCR duplicates and reads with a mapping quality score of less than 10 using Picard's MarkDuplicates and samtools view, respectively (Li et al., 2009), merged BAM files from the same individual, and performed variant calling using GATK HaplotypeCaller (Poplin et al., 2018; Van der Auwera et al., 2013). We identified biallelic SNPs and indels in these regions using GenotypeGVCFs, followed by hard filtering to remove variants with Phred-scaled quality scores < 20 and genotypes with Phred-scaled quality scores < 20 (GATK version 4.6.0.0 (Poplin et al., 2018; Van der Auwera et al., 2013).

### 2. Identification of putative VR genes

We used a set of sequences for VRs from all possible primate genomes including black-capped squirrel monkey, bonobo, chinese tree shrew, common chimpanzee, common marmoset, crab-eating macaque, golden snub-nosed monkey, gray mouse lemur, green monkey, human, northern greater galago, northern white-cheeked gibbon, Philippine tarsier, rhesus macaque, sumatran orangutan, and western gorilla (Moriya-Ito et al., 2018; Hohenbrink et al., 2017; Hunnicutt et al., 2020).

Candidate VR loci were identified by aligning known VR nucleotide sequences from above mentioned species against the white faced capuchin reference genome using BLASTN with default alignment parameters and an initial E-value threshold of $1 \times 10^{-5}$. BLAST hits were subsequently filtered to retain alignments with an E-value $< 1 \times 10^{-10}$, percent identity ≥70%, alignment length ≥200 bp, and bit score ≥100. Filtered alignments were converted to genomic coordinates in BED format and sorted by scaffold and genomic position. To account for fragmented alignments arising from the same genomic region, overlapping or nearby hits separated by ≤1 kb were merged using BEDTools merge utility. During the merging step, the identities of matching query sequences, the maximum bit score, and the minimum E-value for each merged interval were retained. The final output consisted of a non-redundant set of candidate VR loci, including their genomic coordinates, the number of unique query sequences contributing to each locus, the highest alignment bit score, and the most significant E-value, which were used for downstream analyses.

Phylogenetic relationships among VR sequences were inferred using the Maximum Likelihood (ML) method implemented in MEGA. Nucleotide sequences were first aligned using the MUSCLE algorithm with default parameters. The best-fitting nucleotide substitution model was identified using the model selection function in MEGA based on the Bayesian Information Criterion (BIC), which selected the Tamura 3-parameter model with a gamma-distributed rate variation among sites (T92+G) as the optimal model. A Maximum Likelihood phylogenetic tree was then reconstructed under the T92+G model, and branch support was assessed using 1,000 bootstrap replicates. The resulting phylogeny was used to evaluate evolutionary relationships among VR sequences and to identify putative orthologous and paralogous gene groups. Any VR gene for which a name existed in genome assembly, we retained the same name, for other VR genes grouped with these previously named VR genes were named as their paralogs following an alphabetic suffix. For the genes with no names existing within current genome assembly, a name is devised based on orthologous information from the nearest black-capped squirrel monkey genome.

We then extracted the sequences of these VRs to validate the intact/disrupted classifications. We used EMBOSS getorf (Rice et al., 2000) to confirm that the open reading frame of each VR gene was at least 250 amino acids long, the length necessary to form a seven-transmembrane structure required by functional G-protein coupled receptors. We then used DeepTMHMM, a neural network-based approach (Hallgren et al., 2022), to identify open reading frames of 250 amino acids or longer that are capable of making a seven-transmembrane structure. If a VR gene met both criteria, then we considered it intact; otherwise, it was labelled as disrupted. For the identified VR genes with lengths between 500 and 750 nucleotides were labelled as truncated. We note that our analyses are based solely on genomic assessment and our results do not confirm the genes' functional status. For this reason, we use "intact" and "disrupted" gene terminologies rather than "functional" and "pseudogenes."
